# Cerebellar encoding of prior knowledge of temporal statistics

**DOI:** 10.1101/2024.08.19.608550

**Authors:** Julius Koppen, Marit Runge, Lucas Bayones, Ilse Klinkhamer, Devika Narain

## Abstract

We possess the ability to anticipate and preempt occurrences under familiar circumstances, which suggests a reliance on prior experience with regularities in our environment^1–6^, especially when observations become more uncertain. We know little about the neural mechanisms that can learn the probabilities of events in the environment and use this prior experience to guide actions. To examine this, we studied a rudimentary predictive behavior, eyeblink conditioning, and asked whether a simple effector like the eyelid could adapt its movements to varying probabilities of environmental events that reflect different degrees of uncertainty. We found that predictive eyeblink behavior systematically changed almost all its properties according to the temporal statistics of stimulus probability distributions. We also found that the activity of cerebellar Purkinje cells and putative molecular layer interneurons changed concomitantly with temporal statistics of the stimuli and with behavior. Targeted optogenetic perturbation of Purkinje cells during critical time windows severely attenuated the predictive behavior but left reflexive eyeblinks intact. Furthermore, we discovered a novel Purkinje cell complex spike signal coinciding with the onset of the earliest probable time interval in prior distributions with high uncertainty. This signal could not be explained as a motor or sensory correlate and appears to be anticipatory in nature. Theoretical modeling results pointed to a possible synaptic mechanism for how Purkinje cells could encode prior experience of environmental statistics in their activity through the juxtaposition of long-term depression and potentiation dynamics.

## Introduction

Information from the world received by our sense organs is often ambiguous, corrupted by noise, and is inadvertently transformed as it traverses neural circuits, giving rise to uncertain representations. With such uncertainty pervading our sensory observations, we could expect that much of animal behavior relies on inference that incorporates an important source of information, prior knowledge of the environment. This tenet has been formalized using probabilistic frameworks in behavioral and neuroscience disciplines ^3,4,6–8^. Having a form of memory that encodes priors, which reflect the statistics of environmental variables, could be beneficial to survival in many instances, ranging from the average angle of approach of a local predator to the statistics of traffic on daily commutes. We, however, know little about neural circuit mechanisms responsible for learning and distilling the statistics of variables in the world to guide anticipatory behavior.

We can infer from previous work that, consistent with Bayesian theories, prior knowledge can be internalized to warp perception in a manner that improves the precision of various sensorimotor behaviors in humans ^2,8–16^. We also know of cortical representations of such behaviors in sensory ^17–19^ and motor systems ^20^. Recently, Bayesian computations have also been investigated for temporal behaviors, where the activity of neurons in the dorsomedial frontal cortex of monkeys appeared to modulate at the onset of the earliest probable interval in a probabilistic distribution ^21^. The question then arises, if humans^2,14,22^, nonhuman primates ^21^, and rodents ^23^ can learn a variety of tasks with probabilistic stimuli, what are the neural mechanisms by which such probability distributions are acquired and utilized as prior knowledge?

One structure that seems uniquely poised to straddle sensory and motor domains in the context of timing is the cerebellum. The cerebellar cortical machinery has long been implicated in pure timing behaviors^24,25^ alongside various temporally-dependent sensorimotor behaviors^26–38^. Furthermore, several theories propound the cerebellum to be a site for encoding internal models^39–41^ for motor systems^41^, ocular function^42^, vestibular control^43,44^, and inertial perception ^45,46^. Finally, the principal neuron of the cerebellar cortex, the Purkinje cell, exhibits dual firing patterns, low-frequency complex spikes (CSpk), and high-frequency simple spikes (SSpk), which could enable it to synthesize and supervise the learning of a variety of temporal contingencies^47^.

Here, we investigate whether cerebellar Purkinje cells could learn the temporal statistics of intervals sampled from different probability distributions. We ask whether cerebellar cortical populations could give rise to sophisticated predictive eyeblink behaviors that reflect the changing stimulus probabilities of such temporal variables.

### Predictive eyelid behavior modulates with changes in temporal statistics

In the classical version of trace eyeblink conditioning, animals learn a fixed temporal relationship between a conditioned stimulus (light), and an unconditioned stimulus (periocular airpuff), which results in the reflexive closure of the eye (Figure 1a). After repeated pairings, the eyelid exhibits a conditioned response such that the eye closes in anticipation of the trained interval, even in the absence of the airpuff. We refer to this as the predictive response (Figure 1a right). Here, we developed a probabilistic version of trace conditioning, such that the time interval between the light and airpuff on each trial was drawn from different probability distributions or experimentally-defined *prior distributions* (Figure 1b,c).

**Figure 1:**
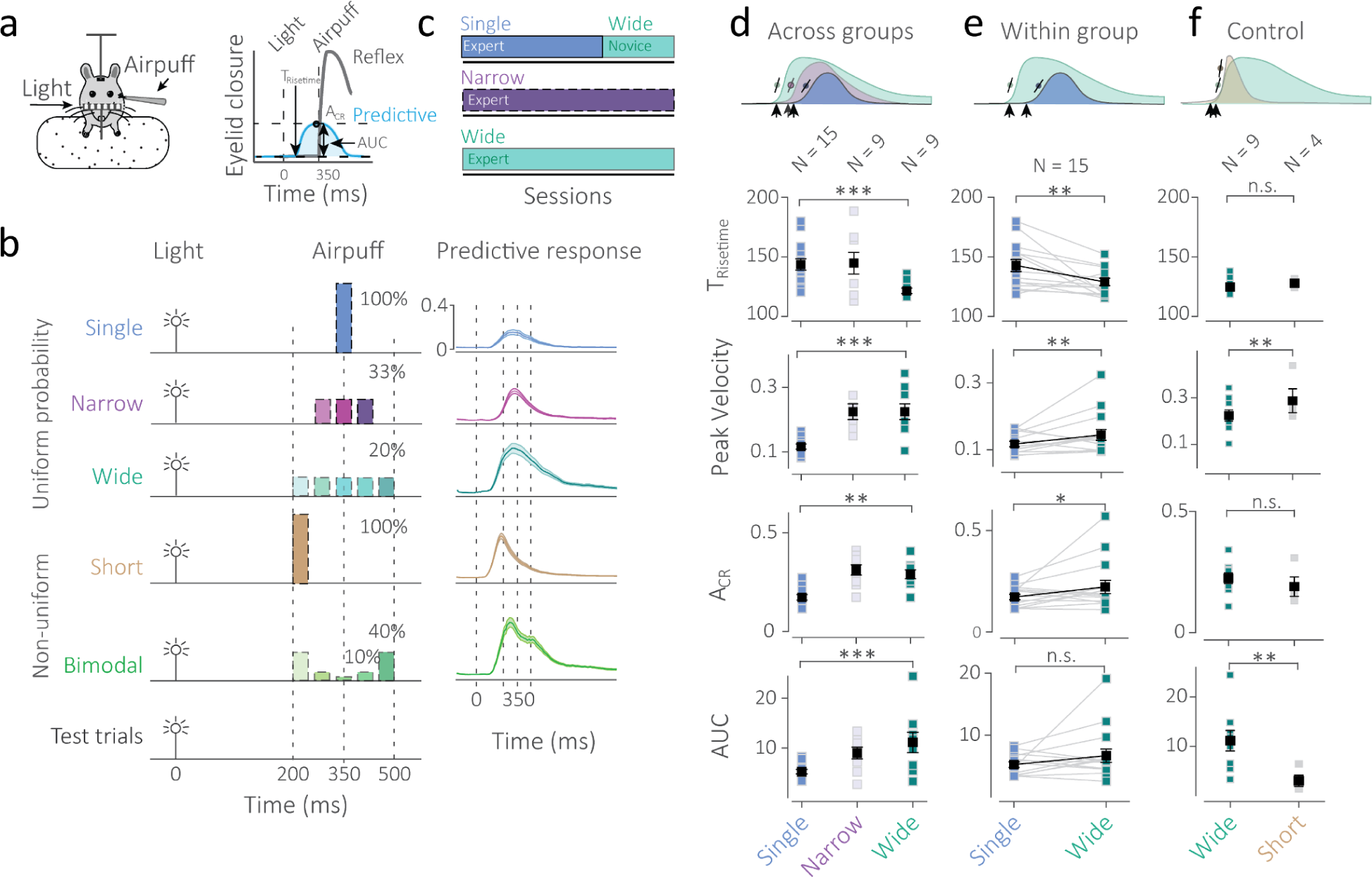
Predictive eyeblink behaviors imprint temporal statistics. a) Left: Head-fixed mice learn temporal delays between a light flash and a periocular airpuff. Right: The airpuff elicits reflexive eye closure (gray) on paired trials but after training, the eye closes preemptively (blue), even in the absence of the airpuff (on test trials), known as a predictive or conditioned response. b) Stimuli are sampled from different probability distributions; *Single* represents a 100% probability of the airpuff on paired trials at 350 ms (blue), *Narrow* represents a uniform probability of 33% (purple), *Wide* represents a uniform probability of 20% on paired trials (teal). The *Short* condition is a control with 100% probability at 200 ms (brown). In the *Bimodal* condition, the stimulus intervals are nonuniform (green). Right: Examples of the average predictive eyeblink response in a subject on test trials (airpuff omitted). c) In addition to expert groups of subjects that were trained throughout on a prior condition, the same subjects were trained on the *Single* and switched to the *Wide* condition (*switch* group). d) Metric comparisons for T_risetime_, peak velocity, A_CR_, and area under the eyeblink response curve (AUC) across the three uniform groups (left), within the *switch* group (middle), and comparison of the *Wide* condition with the *Short* condition group. Black squares represent averages, error bars indicate standard error. * represents p < 0.05, ** p < 0.005 and *** p < 0.0005.

The temporal statistics of these prior distributions varied in the uncertainty of the presented intervals. In the *Single* prior condition, on paired trials, the airpuff always arrived at the same interval, *i.e.*, with a probability of 1 (100%), whereas, intervals in the *Narrow* and *Wide* distributions carried higher uncertainty on paired trials where each was uniformly sampled with a probability of 0.33 (33.3%) or 0.2 (20%), respectively (Figure 1b). We additionally tested stimuli sampled with nonuniform probabilities in a *Bimodal* prior where the shortest and longest intervals were most probable (0.4 or 40% each) and the intermediate intervals were less probable (Figure 1b). Across all these conditions, the mean interval remained constant (350 ms) while the variance, probability mass, and multimodality varied. We also tested a control condition called *Short*, where the probability of a fixed interval of 200 ms was 1 (100%) on paired trials. Mice were either trained throughout on one condition (*expert* mice), or they were trained on the *Single* prior and switched to the *Wide* prior (*switch* mice; Figure 1c).

To examine whether predictive eyeblink metrics altered systematically when the temporal statistics of the stimuli changed, we evaluated blink-related metrics on randomly interspersed test trials (Figure 1b, Supplementary figure 1a), where the airpuff was omitted and only anticipatory behavior could be observed, disentangled from the reflexive eyeblink component caused by the airpuff. These metrics included rise time (T_risetime_), which indicates the earliest time the eye closed predictively, peak velocity of the initial eyelid closure, amplitude (A_CR_) of the predictive response, and area under the curve of the eyeblink trace (AUC) (Figure 1a, right). We found that the metrics of the predictive eyeblink altered systematically as the probability distributions changed statistics. The T_risetime_ was smaller for the *Wide* prior (N = 9), whose earliest probable interval is much shorter in duration compared to *Narrow* (N = 9) and *Single* (N = 15) groups (Figure 1d, F(2,33), = 6.13, p < 0.01) among expert groups. Compared to the *Single* condition, there was a significant increase in peak velocity of eyelid closure for the wider conditions (Figure 1d, F(2,33) = 18.6, p < 0.0001).

As the uncertainty of the prior distributions increased, the probability of the occurrence of each interval also decreased. We hypothesized that this may influence the balance of learning and forgetting due to the presence of the instructive airpuff and the extinction or forgetting that occurs in its absence, respectively. In other words, there may be weaker learning occurring at all intervals for wider priors, leading to a lower amplitude and AUC for wider conditions. To our surprise, the A_CR_ of the predictive eyelid closure increased for the *Narrow* and *Wide* priors compared to the *Single* condition (Figure 1d, F(2,33)= 12.14, p < 0.0001). The area under the eyelid curve (AUC) also increased as the width of the prior distribution increased (Figure 1d, F(2,33) = 8.95, p < 0.001) despite the decrease in the probability of each interval contained.

These patterns also held for the behavior of mice that switched from a *Single* to *Wide* prior condition (N = 15, Figure 1e, Supplementary figure 1b, Supplementary figure 2a-c). Like the expert groups, T_risetime_ decreased (Figure 1e, t(14) = -2.98, p = 0.004), peak velocity increased (Figure 1e, t(14) = 2.06, p = 0.02), and A_CR_ increased (Figure 1e, t(14) = 1.81, p = 0.045) as the mice switched from the *Single* to *Wide* prior. We could not find a significant increase in the AUC for a switch from *Single* to *Wide* (Figure 1e bottom, t(14) = 1.36, p = 0.96), potentially because learning for longer durations may require more training.

An alternative explanation for these results could be that the eyeblink system learns the earliest relevant time interval, making no note of the full temporal distribution or the probabilities involved. To investigate this, we examine the results of a *Short* prior, which, on paired trials, has a 100% probability of occurrence at 200 ms. While the *Short* and *Wide* conditions have the 200 ms interval in common, the probability of its occurrence on paired trials was 1 (100%) for the *Short* condition but only 0.2 (20%) for the *Wide* condition. When we compared these conditions, we found no difference in T_risetime_ (t(11) = 0.79, p = 0.44, Figure 1f) or A_CR_ (t(11) = 0.82, p = 0.42, Figure 1f), however, the peak velocity and area under curve (AUC) were significantly different. The peak velocity was significantly lower for the *Wide* condition (t(11) = 2.38, p = 0.02, Figure 1f) and the AUC was larger for the *Wide* condition as well (t(11) = 2.48, p = 0.015, Figure 1f), both suggesting that learning is weaker for the 200 ms interval in the *Wide* condition and that the eyeblink kinematics slow down in their later phase to, potentially, accommodate the longer range of the *Wide* distribution.

Further evidence that the probabilities of the full distribution are taken into account in the predictive eyeblink can be seen in mice trained on a *Bimodal* prior condition with a nonuniform probability distribution (Figure 1b, Supplementary figure 1a). We found that predictive eyeblinks exhibited multimodality, displaying two peaks in the average conditioned eyelid response at times that corresponded well with the durations of higher probability in the *Bimodal* prior condition (Figure 1b, Supplementary figure 1a). Overall, we found that the predictive eyelid responses systematically altered their timing, size, width, and kinematic properties according to increasing uncertainty in the probability distribution of the temporal stimulus in trace eyeblink conditioning.

### Cerebellar cortical activity changes concomitantly with temporal statistics and behavior

We asked whether neurons in the cerebellar cortex would concomitantly change the properties of their modulation with temporal statistics of the stimuli. Using an anatomical grid-in-chamber setup, we performed acute large-scale electrophysiological recordings of neurons in lobules IV/V and Simplex from the molecular and Purkinje layers (see methods, Figure 2a,b) of the cerebellar cortex. We quantified probe locations based on histological analysis of tracks by registering them to the Allen Brain Atlas Common Coordinate Framework^48^ (Supplementary figure 3a-c). To understand the functional nature of the neural responses, we used recently developed statistical approaches^49^ to determine the overall modulation of neurons and to functionally classify neurons as responsive to the interval (I) epoch (following the light stimulus to the range of the prior) or the airpuff epoch (A) (Figure 2c, Supplementary figure 4). Subsequently, using physiological features such as baseline rates, complex spike polarities, and simple spike-complex spike cross-correlograms, we identified Purkinje cells (or putative Purkinje cells, Figure 2b,d, see methods), and putative molecular layer interneurons (Figure 2d, Supplementary figure 5a,b).

**Figure 2:**
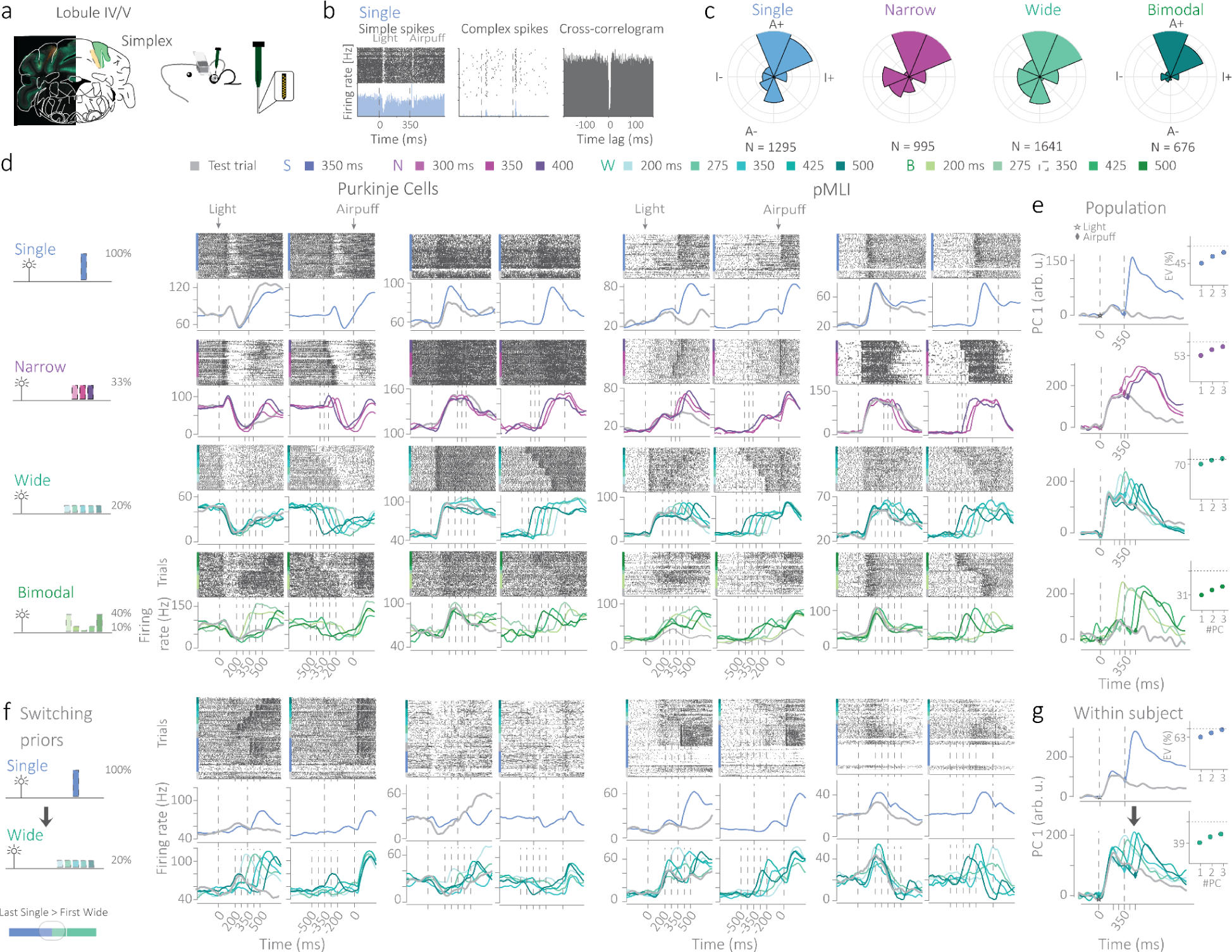
Cerebellar cortical activity encodes temporal statistics. a) Extracellular recordings were made from the molecular and Purkinje layers of Lobule IV/V and Simplex of the cerebellar cortex. b) Purkinje cells were identified by cross-correlations between simple and complex spike activity. c) Functional classification of all statistically modulating neurons recorded in each expert condition. ‘I’ represents the interval epoch and ‘A’ represents alignment to the airpuff. + and - symbols represent the facilitation and suppression of activity in these epochs, respectively. d) Examples of rasters and firing rates of Purkinje cells and putative molecular layer interneurons (MLIs) for different prior conditions (*Single, Narrow, Wide*, and *Bimodal*). Each neuron’s activity is aligned to the light and to the airpuff. e) Representative examples of the first principal component (PC) of a neural population for a single subject (average variance explained by PC1: 43%) for different prior conditions. f) Examples of Purkinje cell and putative MLIs recorded as the subject switched from the *Single* to the *Wide* condition for the first time. g) Primary principal component over time capturing dynamics of cerebellar cortical activity before and after the switch from *Single* to *Wide* in the same subject.

We found a large number of statistically-significantly modulating neurons in expert mice for the *Single* (N_mice_ = 10, N_neurons_ = 2819, Figure 2c,d, Supplementary figure 4, 5a,b, Supplementary table 2), *Narrow* (N_mice_ = 6, N_neurons_ = 3295, Figure 2c,d, Supplementary figure 4, 5a,b, Supplementary table 2), and *Wide* (N_mice_ = 8, N_neurons_ = 2339, Figure 2c,d, Supplementary figure 4, 5a,b, Supplementary table 2) conditions, and also in the *Single-Wide switch* condition (Switch, N_mice_ = 9, N_neurons_ = 5271, Figure 2f, Supplementary figure 5c-d, Supplementary table 2). Furthermore, we recorded modulating cerebellar cortical neurons for the non-uniform *Bimodal* prior condition (*Bimodal*: N_mice_ = 3, N_neurons_ = 1791, Figure 2c,d, Supplementary figure 4, 5a,b, Supplementary table 2). Functional classification of each of these neural populations revealed patterns of heterogeneity in activity profiles (Figure 2c) that were consistent with previous findings^50^.

To examine if we could observe prior-related modulation at the level of population dynamics, we performed a principal component analysis on the neural data for each expert mouse, and in the case of *switch* mice, for each neural population in each condition before and after the switch. We found that the largest principal component (PC), which on average explained approximately 43% of variance (see Table 2) in the neural population, resembled the average behavior of mice for test and paired trials (Figure 2e, Supplementary figure 6a). Both for the *expert* and *switch* mice, the largest principal component also showed the same patterns of changes as observed in the behavioral data, i.e increase in AUC with increase in prior width, earlier T_risetime_ and faster peak velocity (Figure 2e,g, Supplementary figure 6a,c). It also exhibited more complex patterns like dual peaks for the *Bimodal* condition (Figure 2e, Supplementary figure 6b).

To quantify whether cerebellar cortical responses systematically changed with temporal statistics, we leveraged trial-by-trial decoding using a machine learning inference technique known as LFADS^51^ (Latent Factor Analysis via Dynamical Systems). LFADS infers the initial conditions and dynamics underlying spiking patterns in neuronal data and generates predictions of firing rates of single neurons on a trial-by-trial basis. We show that LFADS successfully recapitulated neural activity profiles for a large proportion of neurons (Figure 3a,b, Supplementary figure 7a,b). Metrics extracted on a trial-by-trial basis from the neural activity were found to correlate with the behavioral metrics (Figure 3c, Supplementary figure 8a-c). They also recapitulated the same patterns observed for metric changes for T_risetime_, peak velocity, and A_CR_ in all mice on which the LFADS analysis was successfully performed (Figure 3c, Statistics reported in table 1). The AUC metric was also found to be significant in most cases (Figure 3c, Statistics reported in table 1). In summary, we show both qualitatively (principal component analysis, Figure 2e,g, Supplementary figure 6a-c) and quantitatively (trial-by-trial decoding of neural activity, Figure 3c, Supplementary figure 7a,b; 8a-c) that concomitant with behavioral changes, cerebellar cortical activity systematically modulated based on the changing uncertainty of prior statistics.

**Figure 3:**
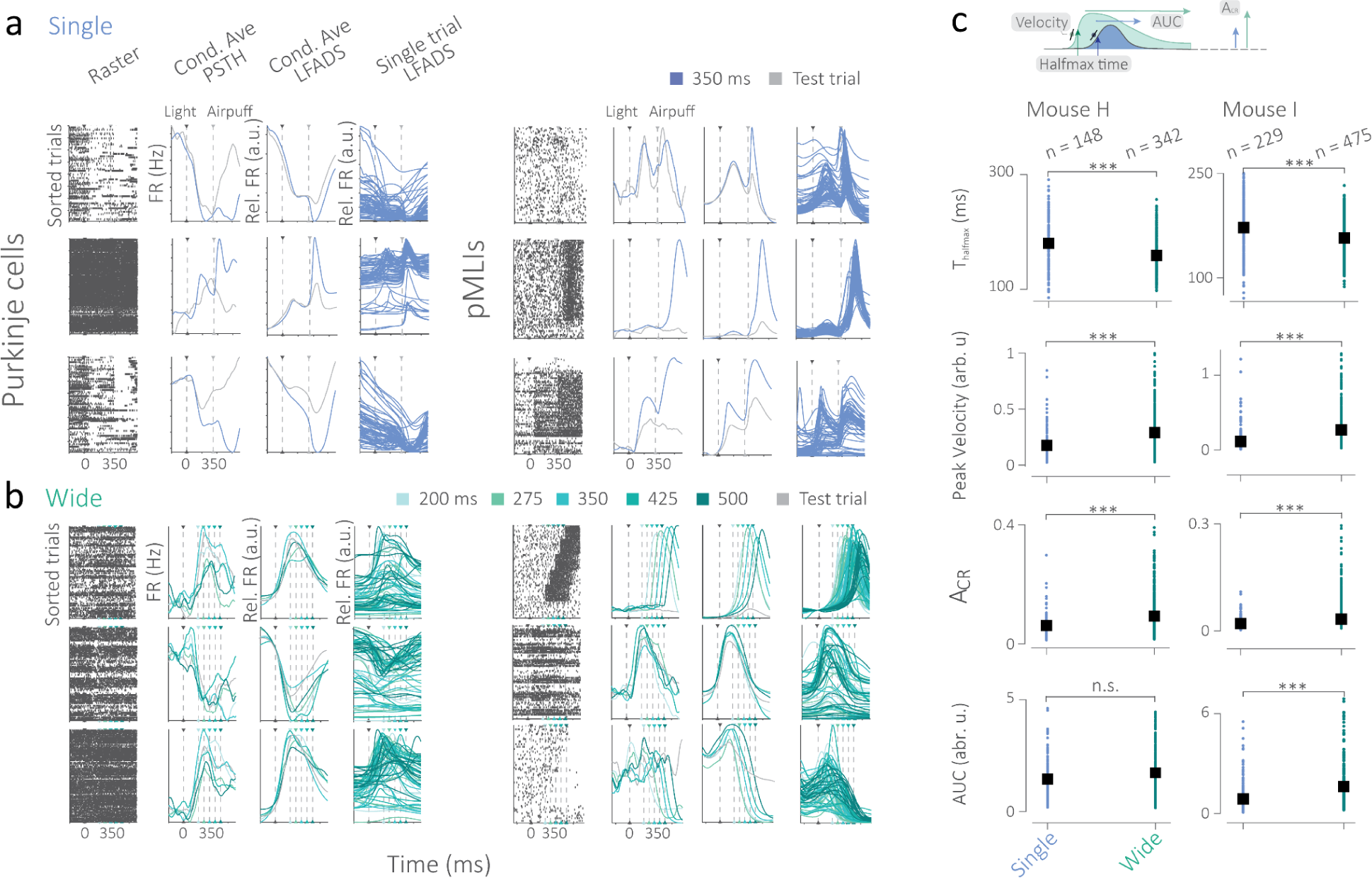
Trial-by-trial decoding of metrics from neural data recapitulates behavioral patterns. a) Left to right: Spike times organized trial-wise (raster), conditioned average firing rates, historically known as peri-stimulus time histograms or PSTH, conditioned averaged LFADS-inferred activity, LFADS inference of activity on individual trials in the *Single* condition. Left: examples of Purkinje cells and putative Purkinje cells, Right: examples of putative molecular layer interneurons. b) Same as a) but for the *Wide* condition. c) Metrics extracted from LFADS-inferred trial-by-trial estimates of neural activity of the same subjects (N = 3) as they switched from *Single* to *Wide*, recapitulate the behavioral decrease in halfmax time, increase in peak velocity, A_CR_ and AUC. Black squares represent averages, error bars represent standard error. *** represents p < 0.0005. Full statistics in table 1.

**Table 1:** T-statistic and p-values of a two-sample t-test for the LFADS behavioral metric comparisons for neural and behavioral data from the Single and Wide conditions.

|  | Mouse H |  | Mouse I |  | Mouse D |  |
| --- | --- | --- | --- | --- | --- | --- |
| Halfmax time | t(488) = -5.58 | P < 0.0001 | t(702) = -6.03 | P < 0.0001 | t(767) = -16.3 | P < 0.0001 |
| Peak Velocity | t(488) = 3.7 | P < 0.0001 | t(702) = 6.68 | P < 0.0001 | t(767) = 8.92 | P < 0.0001 |
| A<sub>CR</sub> | t(488) = 6.05 | P < 0.0001 | t(702) = 7.34 | P < 0.0001 | t(767) = 8.92 | P < 0.0001 |
| AUC | t(488) = 1.88 | P = 0.06 | t(702) = 5.17 | P < 0.0001 | t(767) = 3.41 | P < 0.001 |

### A novel Purkinje cell complex spike signal encoding the onset of prior distributions

Previous work on classical delay eyeblink conditioning has reported the occurrence of Purkinje cell complex spike activity following the light^28,52^ (CSpk_light_) and following a periocular airpuff^53,54^ (CSpk_airpuff_). We found both of these principal types of functional complex spike signals, CSpk_light_ (N*_Single_* = 231, N*_Wide_* = 360) and CSpk_airpuff_ (N*_Single_* = 289, N*_Wide_*= 288) in the *Single* and *Wide* prior conditions (Figure 4a-d, Supplementary figure 9). These complex spikes were characterized using a recent statistical method^49^ and based on this we report their prevalence in our neural data (Figure 4e). We characterized their width and latency compared to the onset of the two sensory stimuli they are associated with *i.e.*, the light and airpuff, respectively (Figure 4f,g).

**Figure 4:**
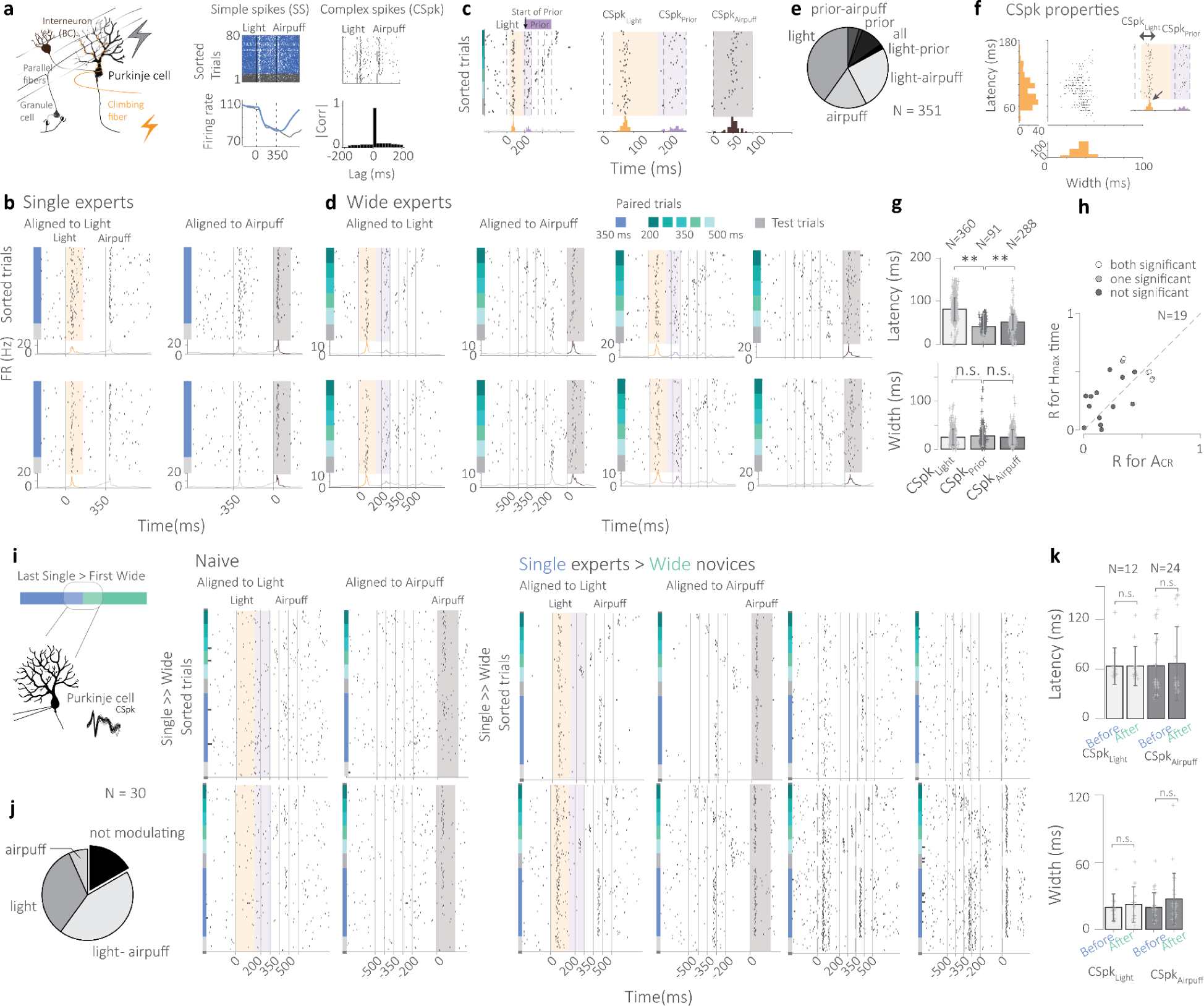
A complex spike signal that marks the onset of high uncertainty prior distributions. a) Purkinje cells exhibit high frequency simple spike activity and low-frequency complex spike activity, which when cross-correlated reveal a 20 ms suppression of simple spikes by complex spikes. b) In the *Single* condition, earlier findings of a light-related complex spike (CSpk_light_, in orange) and an airpuff-related CSpk_airpuff_, in gray, are recapitulated. c-d) In *Wide experts*, in addition to the CSpk_light_ and CSpk_airpuff_, a prior-related complex spike CSpk_prior_ is found, which is time-locked to the onset of the earliest probable interval of the *Wide* prior (in purple). e) Relative proportions of the three functional types of complex spikes found in *Wide experts*. f-g) The width and latency of the CSpk_light,_ CSpk_airpuff_, and CSpk_prior_ are calculated from the onset of the light, airpuff and first interval of the prior, respectively. h) Correlations of the time of the CSpk_prior_ with behavioral metrics such as T_halfmax_ and A_CR_. Non-significant correlations are shown in black and significant correlations in white or light gray. i) Purkinje cell complex spikes were recorded in Naive subjects and in *switch* mice as they transition from *Single* to *Wide*. j) Percentage modulation and the functional categories of complex spikes identified in the mice that switched from *Single* to *Wide*. No CSpk_prior_ were found. k) Latency and width for the CSpk_light,_ CSpk_airpuff_ before and after the switch. Crosses represent individual values, bars represent averages. Error bars represent standard error.

In mice trained on the *Wide* condition, we encountered an unexpected complex spike signal in Purkinje cells that appeared to be coincident with the onset of the earliest probable interval of the prior distribution. We refer to this as the prior-related complex spike (CSpk_prior_, Figure 4c,d, Supplementary Figure 10a,b). The prevalence of statistically-significant CSpk_prior_ among all complex spikes recorded in *Wide experts* was determined to be 21% (Figure 4e, N = 91). Unlike the CSpk_light_ and CSpk_airpuff_, there is no external sensory event that is tied to the vicinity of the CSpk_prior_ in a vast majority of trials. If we, however, compute its latency from the onset of the earliest probable interval in the prior (200 ms for the *Wide* prior), we find this latency to be smaller than that obtained for the CSpk_light_ (t(449) = 13.4, p = 1.7·10^-34^) and CSpk_airpuff_ (t(377) = -2.94, p = 0.0034) relative to the light and airpuff, respectively (Figure 4g, Supplementary figure 11a,b). The precision (or width) of all three types of complex spikes could not be distinguished (Figure 4g, Supplementary figure 11a,b, Prior-Light: t(449)=1.41, p = 0.159, Prior-Airpuff: t(377) = -0.65, p = 0.52). The trial-by-trial time of onset of the CSpk_prior_ does not appear to be highly correlated with behavioral metrics of the eyeblink such as T_risetime_ or A_CR_ for Purkinje cells that had a sufficient number of behavioral trials and prior-related complex spikes (Figure 4h, Supplementary figure 12).

We performed a control experiment to rule out whether the CSpk_prior_ could represent a passive stimulus-related phenomenon that may be an intrinsic biophysical property and not an acquired or learned feature. We recorded from Purkinje cells when the mouse first switched from the *Single* to *Wide* condition and examine the properties of the complex spikes (N_neurons_ = 30, Figure 4i). We did this for both naive groups of mice (N = 5) and a subset of *Single experts* who would switch to the *Wide* prior (N = 5). We found that the CSpk_airpuff_ was present in naive and expert mice (N_neurons_ = 49, 67%). On the other hand, the CSpk_light_ was present mainly in the expert group (N_neurons_ = 12, 40%). The properties of the CSpk_light_ (width: t(11) = -0.043, p = 0.97, latency: t(11) = -0.304, p = 0.77, Figure 4k) and CSpk_airpuf_ (width: t(23) = -1.290, p = 0.21, latency: t(23) = -0.618, p = 0.54, Figure 4k) did not change before and after the occurrence of the switch. More importantly, we could not find the occurrence of any CSpk_prior_ in the naive, *Single* expert, or *Wide* novice groups.

These results show that the CSpk_prior_ does not appear to be present at the first instance that the cerebellar cortex encounters high uncertainty temporal distributions but is likely to be prevalent in mice with prolonged exposure to high uncertainty distributions. This indicates that the CSpk_prior_ may arise as a consequence of learning within the cerebello-olivary loop and may predict the onset of the prior. We do not find strong evidence of correlation of the time of CSpk_prior_ with motor aspects of the behavior. Furthermore, the latency of the CSpk_prior_ with respect to the onset of the prior is significantly lower than the latencies exhibited by the CSpk_light_ and the CSpk_airpuff_ with respect to the light and airpuff, respectively.

### Optogenetic perturbation reveals cerebellar cortical involvement in prior-related activity and behavior

To examine the role of the Purkinje cell in eliciting prior-related behaviors, we turn to existing theories of suppression of Purkinje cell simple spike activity as the primary mechanism that drives conditioned eyeblinks ^28,55^. Since the area we investigate is broader than that proposed in earlier works ^28,55^, we confirm the involvement of Purkinje cells in Lobule IV/V and Simplex by an optogenetic strategy that directly targets Purkinje cell suppression in these areas during the prior-related interval window, which we predict should lead to a decrement in predictive eyelid closure but should leave the reflexive component of the eyelid movement, elicited by the airpuff, intact. We expressed Channelrhodopsin2 (ChR2) in cerebellar Purkinje cells of mice (PCP2-Ai32-eYFP^56^), implanted a tapered optic fiber in lobule simplex, and simultaneously recorded cerebellar cortical neurons (Figure 5a, Supplementary figure 13a,b). We administered calibrated low-intensity pulsed stimulation on each trial just preceding the onset of the first interval of the prior distribution at random on 40% of trials within a session (opto trials). We refer to the remaining trials as control trials. On opto trials, we observed an elimination or severe attenuation of the predictive component of the eyeblink (Figure 5b,c). In all cases, the reflexive component of the same motor behavior remained fully intact (Figure 5c), which is believed to be influenced by a different cerebellar ^57^ and reflex pathway^58^. Accordingly, when we compared the control and opto trials, we found that the CR percentage (t(4) = -2.3, p < 0.05), A_CR_ (t(4) = -1.98, p < 0.05) and AUC (t(4) = -2.1, p < 0.05) all significantly decreased (Figure 5d).

**Figure 5:**
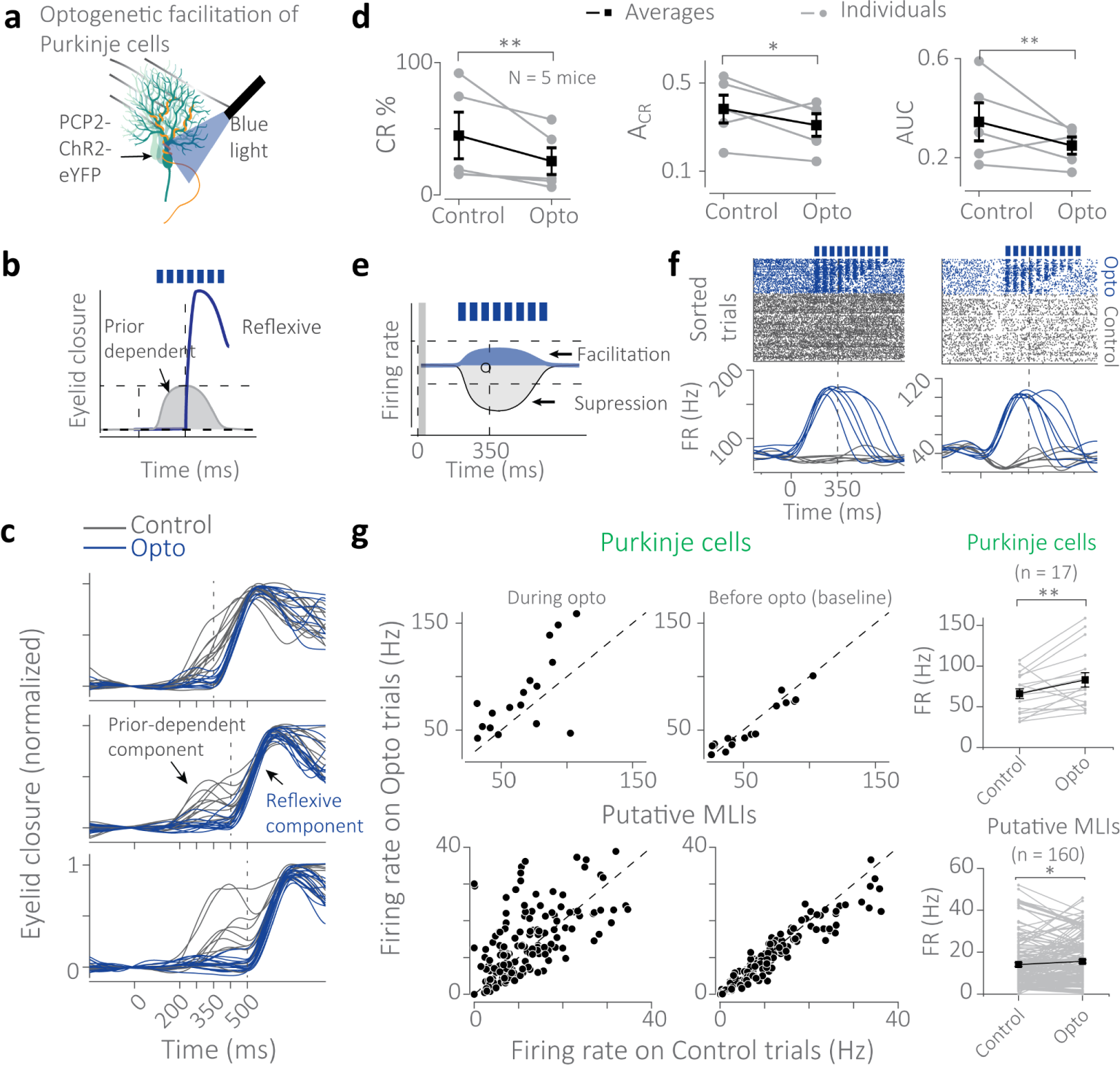
Optogenetic silencing of the prior-related predictive component of the eyeblink. a,b) Optogenetic strategy to probe the role of suppression of Purkinje cell activity during the predictive behavioral response: Mutant mice expressing Channelrhodopsin2 in Purkinje cells (PCP2-Cre-ChR2-eYFP) were trained on the *Wide* prior condition. Electrophysiological recordings were performed as optogenetic perturbation with blue light was administered at random on 40% of trials within a session. c) Individual behavioral traces for different time intervals comparing trials with optogenetic perturbation (dark blue) vs. controls (gray) within the same session on paired trials where, for normal conditions, both prior-dependent and reflexive components are expected to be present. d) Behavioral summary for A_CR_, CR percentage, and AUC comparing optogenetic and control trials within sessions of the same mouse. Black circles and lines indicate averages, whereas gray circles and lines indicate individuals. Error bars represent standard error. e) Precise optogenetics to wash out suppression dynamics during prior-related behavior. f) Rasters and firing rates of Purkinje cells during optogenetic sessions revealed the successful facilitation of these cells while remaining in the natural activity regime of Purkinje cells. g) Top: Majority of Purkinje cells (n = 17) and putative MLIs (n = 160) recorded exhibited a significant increase in firing rate during optogenetics trials compared to control trials (left) only during the optogenetic epoch and not during the baseline epoch (middle). Right: In general, both Purkinje cells and putative MLIs registered an increase in firing rate during optogenetic trials compared to that during control trials. Black lines and circles represent averages. Error bars are standard error.

To ascertain that the optogenetic stimulation was correctly and specifically administered, we analyzed our neural recordings during these sessions (Figure 5e-f, Supplementary figure 13b). The firing rate of identified Purkinje cells was significantly higher during stimulation than that during the baseline epoch of the same cell in the same session (t(16) = 2.5, p = 0.01, Figure 5g), showing no long-term disruption of neural activity. For putative molecular layer interneurons, there was greater variability in the outcome but these also registered a significant increase in overall activity compared to their baseline epochs (t(159) = 2.2, p = 0.02, Figure 5g). These results show that precisely timed facilitatory perturbation during the range of the prior in simplex lobule Purkinje cells severely and reversibly impairs the prior-dependent component of the learned eyeblink response while leaving the reflexive component of the same behavior intact.

### A computational cerebellar mechanism for the acquisition of temporal statistics

We built a computational model that utilizes earlier proposals^50,59–64^, to advance a novel link between continuous-time eyeblink behavior and the formation of prior-dependent memory of timing in cerebellar cortical circuits. Earlier proposals in cerebellar cortical theory have focussed on long-term plasticity mechanisms activated as a function of convergent sensory signals arising at the principal cerebellar cortical neuron, the Purkinje cell. During a trial, the model assumes that the light cue activates a cascade of granule cells (GC), giving rise to a temporal basis (Figure 6a), which appears to be consistent with recent evidence^65–68^ and whose shape accommodates scalar variability in timing^64^. The computational model hypothesizes that each of these hundred thousands of granule cells make synapses with Purkinje cells, funneling in potentially temporally heterogeneous but repeatable inputs time-locked to the onset of the light. After the interval elapses and the airpuff arrives, the proprioceptive response is assumed to activate the inferior olive resulting in the generation of an airpuff-related complex spike in Purkinje cells. The conjunctive activation of GCs and climbing fibers is generally believed to result in long-term depression (LTD) of the subset of GC-Purkinje cell synapses active at the time close to the airpuff. In addition to this, the model employs a long-term potentiation mechanism in the absence of climbing fiber signals that seeks to return the system to homeostasis (Figure 6a).

**Figure 6:**
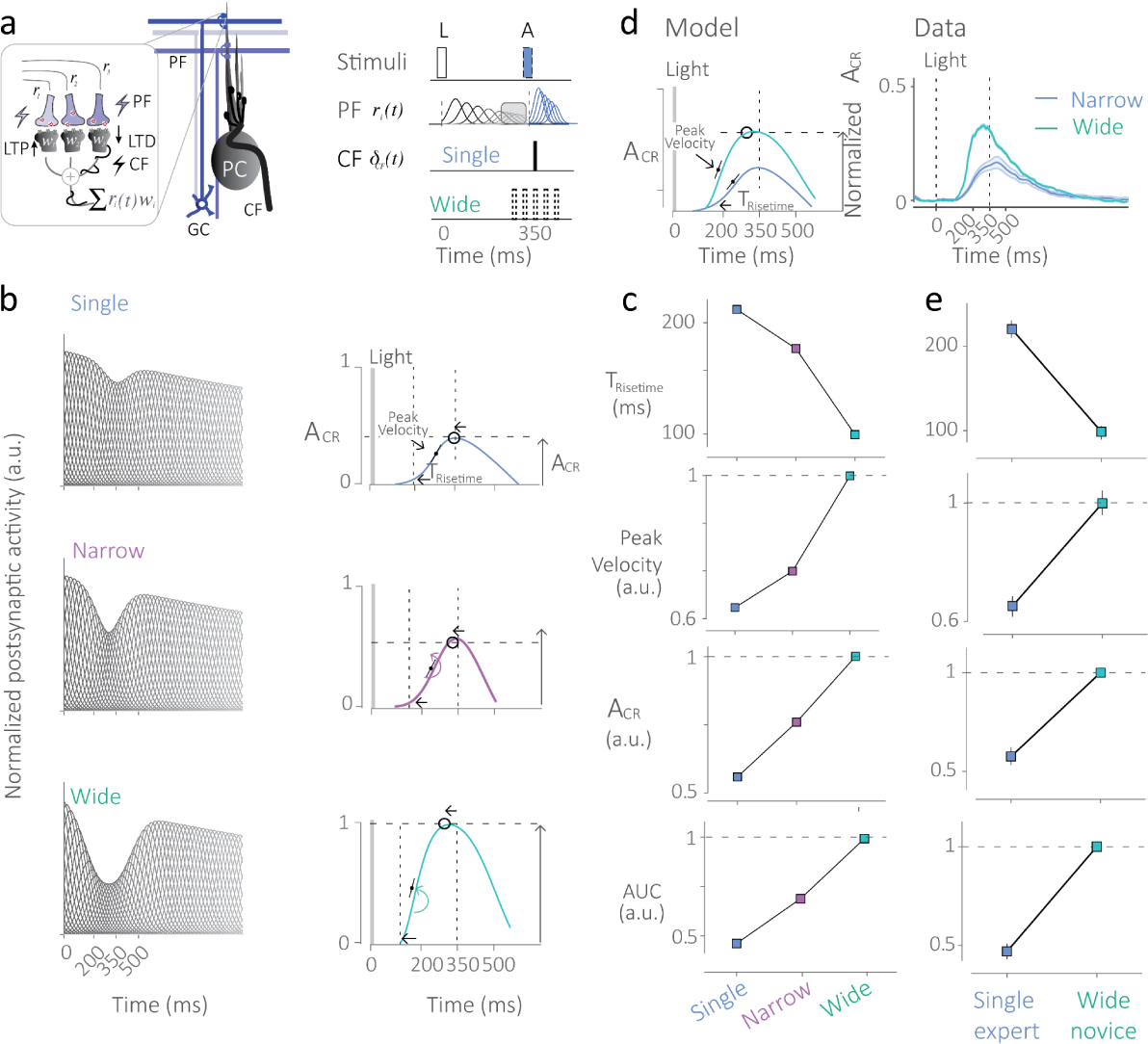
A cerebellar cortical mechanism for learning temporal statistics. a) Cerebellar microcircuitry consisting of inputs to the Purkinje cell via granule cell (GC) parallel fibers (PF) and olivary climbing fibers (CFs). The model assumes that conjunctive activation of the GC and CF pathways leads to long-term depression (LTD) and activation of PFs alone leads to long-term potentiation (LTP) of granule cell to Purkinje cell synapses. Purkinje cell activity above baseline is assumed to be a weighted sum of PF-related inputs. b) Model predictions for the postsynaptic activity after learning of temporal statistics. These results ensue from the gating of granule cell-Purkinje synaptic weights leading to a decrease in net input activity that imprints the properties of the temporal distribution. Right: Averaging of these inputs and downstream linear combinations lead to model predictions for behavior for each prior condition. c) The decrease in T_risetime_, increase in peak velocity, A_CR_, and AUC are recapitulated for the *Single* (blue), *Narrow* (purple), and *Wide* (teal) conditions. d) Model predictions and example of the behavior of a representative subject that switched from *Single* to *Wide*. e) The model recapitulates the behavior observed for behavioral patterns following within subject switches from the *Single* to *Wide* condition.

Since, in the probabilistic version of trace conditioning, the airpuff arrives at different times and with different probabilities, in the model, the rate of learning (LTD) and forgetting (LTP) is influenced by the probabilities of the intervals, *i.e.* the varied timing of CSpk_airpuff._ This results in the selective gating of incoming granule cell inputs around the relevant intervals, whose synaptic weights mirror the probability of occurrence of the intervals of the distribution (Figure 6b). When these inputs were averaged over time and the net Purkinje cell activity was linearly transformed for downstream areas to generate a motor output, we found that behavioral predictions matched the increase in amplitude, decrease in rise time, increase in peak velocity, and area under the curve, as observed in mice with the increased uncertainty in the prior from *Single* to *Narrow* and from *Narrow* to *Wide* (Figure 6c). Furthermore, when we examine predictions for the within mice switch from *Single* to *Wide* prior (Figure 6d right), we find the same decrease in rise time, increase in peak velocity, amplitude, and AUC (Figure 6d-e).

## Discussion

Most of us remain blithely unaware that our eyes blink in anticipation of an imminent environmental perturbation, such as a tennis ball gone awry or an insect on an apparent collision course with us. Even if we were partly aware of these feats, few would suspect that the eyeblink system could keep track of the probabilities of events in our world. Here we demonstrate that preemptive eyeblink profiles inculcate our prior experience with probability distributions of temporal stimuli.

We show that the activity of cerebellar Purkinje cells encodes these statistics in their simple spike activity and that their complex spike activity can signal the onset of prior distributions. These results show that prior distributions are internalized in Purkinje cell signaling both in its high and low frequency signals.

When we optogenetically perturb Purkinje cell simple spike activity during critical prior-related windows, the predictive eyeblink response is extinguished from behavior, suggesting a relationship between the expression of these temporal statistics in Purkinje cells with those expressed in predictive eyelid behavior.

The prior-related complex spike CSpk_prior_ is a surprising feature of Purkinje cells that only seems to appear after sufficient exposure to the high uncertainty distribution. The CSpk_prior_ has no immediate sensory events in its vicinity, does not correlate well with the motor output but it predicts the expected start of the prior better than the attributes of the CSpk_airpuff_ or the CSpk_light_ do for their respective sensory stimuli. <u>Here we propose that the combination of a higher magnitude of granule cell input at earlier</u> <u>durations alongside additional learning ensuing from the CSpk_prior_ may explain why we see higher</u> <u>amplitudes of learning for early intervals in the higher uncertainty prior conditions.</u>

Earlier work has uncovered cerebral cortical neurons that begin to modulate in response to the onset of the stimulus distributions^21^. These areas are disynaptically connected with the cerebellum and one could hypothesize that the olivary-cerebellar system may generate a predictive signal marking the onset of high uncertainty distributions that may be utilized by downstream areas to carry out complex behaviors.

To understand underlying mechanisms behind this phenomenon, we propose a simple model, based on previous works^50,62–64,69^, that uses the juxtaposition of long-term depression and potentiation to gate synaptic activity of granule cell-Purkinje cell synapses. The probabilistic nature of the stimuli influences the balance of learning and forgetting, which also interacts with the native strength of the hypothesized granule cell basis set, evidence for which is recently gathering momentum^65^. However, there are many aspects of the behavior that this simple model cannot account for. For instance, it does not directly account for the heterogeneity in neural activity observed, which has been addressed by other models^50^. Nor does it account for potential plasticity and tuning of the granule layer through synaptic diversity, which has recently been proposed^70^.

Previous reports in human behavior indicate that humans find it difficult to accurately learn complex priors with multimodality or kurtosis, and behavior for only simpler priors is likely to be consistent with Bayesian inference models^14^. Our results and previous work^14,71^ suggest that eyeblink behavior in rodents and models of cerebellar cortical circuits may explain this observation. The nature of the granule cell basis set and its interaction with Purkinje cell synapses may lead to an inadvertent low-pass filtering that may disadvantage the learning of temporal priors with high kurtosis or other second order statistics.

Despite various demonstrations of how human and nonhuman animal behavior can be consistent with Bayesian theories, skepticism about the perspective persists^72^ and even its proponents advocate a teleological narrative rather than exploring the potential origins of the phenomenon in neural circuits^73^. The result that prior knowledge of the environment can reshape even primitive conditioned behaviors may bolster the waning belief that mammalian behavior is continually optimized by the statistical regularities of our world.

## Methods

### Mice and surgical setup

This study utilized 52 mice (Postnatal age > 60 days, 21 female). C57BL/6 mice were trained as *experts* in groups for five prior conditions: *Single* (N = 16), *Narrow* (N = 9), *Wide* (N = 9), *Short* ( N = 4), and *Bimodal* (N = 3). A subset of *Single experts* were switched to the *Wide* condition and called *switch* mice (N = 15). Naive mice (N = 5) were also used as control subjects for electrophysiological recordings. Six L7-cre (BAC-Pcp2-IRES-Cre^74^) crossed to Ai32 (Rosa26-LSL-ChR2-eYFP^56^, JAX) were trained on the *Wide* condition, of these, five mice were used for optogenetics (N = 5). All procedures were performed in accordance with protocols approved by the animal care and use committees (IvD) at Erasmus Medical Center. Mice were housed in a 12:12 light:dark cycle and were tested in the light phase. No restriction was placed on food. Water dispensation was maintained throughout and body weight was monitored. All surgical procedures were carried out aseptically under a mixture of 3% isoflurane in 1 L/min oxygen anesthesia. Post-operative analgesia management was enabled by administering Buprenorphine HCl (0.1 mg/kg) and Carprofen (5 mg/kg). Mice were monitored and treated for 3 days post-surgery before the continuation of experiments. Between postnatal age P60 and P80, a semi-magnetized pedestal was installed on the skull to enable head fixation during behavioral training. After stabilization of behavioral metrics on the *Single* condition (Supplementary figure 2), a custom-designed thermoplastic recording chamber was installed on a cerebellar craniotomy centered at AP -6.25 mm, ML -2.25 mm from Bregma, and 3 mm in diameter. The chamber was affixed using an adhesive (Optibond, Kerr corporation, USA) and dental cement (Charisma, Kulzer, USA). After implantation, the chamber of each mouse was cleaned daily with saline and a dura-cleaning tool and disinfected with low concentrations of ethanol to maintain the hygiene of the dura and surroundings. All surgical installations on the skull were aligned using 2D line level. During recording sessions, a custom-made plexiglass grid was installed into the chamber to ensure systematic anatomical access to cerebellar structures and to provide stable housing for the electrode. In optogenetics experiments, the tapered optic fiber (Sigma fiber, Optogenix, Italy) was implanted through an adjacent grid location at an angle of 7 degrees.

### Task, Experimental setup, design and training

Mice were head-fixed to a post with a semi-magnetized pedestal attached to their skull and were able to comfortably rest or move freely on a self-initiating treadmill with low forward resistance but adequate textured grip. A high-speed infra-red camera (Basler Ace aca1300, Basler, Germany) was trained on the eye of the animal to record movements. Our setup was designed based on earlier proposals for similar tasks^75^. Posterior whiskers were trimmed to minimize interference with eyeblink detection. A custom-made device delivered a peri-ocular airpuff representing the unconditioned stimulus using an air-pressurized drive triggered by a 5V pulse. A semicircular array of white LEDs was used to deliver the light representing the conditioned stimulus. This delivery system was built to ensure homogenous and adequate bilateral visual input to the mouse (to ensure equivalent bilateral activation of the pontine nuclei). The camera recording and stimulus delivery system were integrated using custom drivers and code in Objective C (Cocoa framework XCode, Apple, Cupertino, USA) and Matlab 2020a (Matlab, Natick, USA). Pulse pal (Sanworks, USA) was used for regulating stimulus delivery.

Each experimental training session lasted for 80 trials, with an inter-trial interval sampled from a discretized truncated exponential function (tau = 4s). Mice were given an expert status when the CR percentage values saturated to or exceeded a threshold of approximately 40% (Supplementary figure 2c). After performance metrics stabilized, mice underwent a surgical craniotomy and chamber placement. After recovery, expert mice continued training on the same condition. For *Single experts*, after several electrophysiological recording sessions with the *Single* condition, behavior was switched to the *Wide* condition. After discontinuing electrophysiological recordings, the mouse continued with behavioral training on the *Wide* condition (Supplementary figure 2c).

On each trial, the LEDs (Light) were active for 70 ms, followed by the interstimulus interval (ISI), after which the airpuff was administered for 70 ms, resulting in reflexive eye closure. The ISI was determined based on the prior condition. For the *Single* prior, the ISI was 350 ms, for the *Narrow*, the ISI was uniformly sampled from a discrete distribution: [300, 350, 400] ms. For the *Wide* prior, it was also sampled with uniform probability from a discrete distribution: [200, 275, 350, 425, 500] ms. The *Short* prior used an ISI of 200 ms. In the *Bimodal* prior, there was a 40% probability of sampling the first and fifth interval and a 10% probability of sampling the second and fourth interval from the distribution [200, 275, 350, 425, 500]. All conditions contained randomly interspersed test trials where the airpuff was omitted and these trials did not influence the probability calculations on paired trials. At the start of each session, two airpuff-only trials were used to compute and calibrate the baseline eye closure by calculating the change in pixel value during each frame in the region of interest. This value was used to normalize subsequent eyeblink responses. Mice were monitored at all times during training and were given a time-out if squinting or extended eye closure was detected.

### Quantification of eyeblink metrics

Eyeblink responses within the session were normalized to the reflexive response to the airpuff, which was recorded in absence of the light stimulus on airpuff-only trials (administered just before the start of the session) and on paired trials. A conditioned response was detected if the eyeblink trace velocity increased beyond threshold and the amplitude exceeded thrice the baseline standard deviation before the light presentation across all trials within a session. 1) CR percentage was computed as the number of trials where a CR was detected against the total number of trials within the session. 2) CR amplitude was computed as the maximum eye closure of the predictive component after light onset. 3) Rise time was computed as the time when the derivative of the eyelid trace exceeded twice the standard deviation of trace across trials within the session. 4) AUC was computed as the sum of the eyeblink trace on test trials within 600 ms of light presentation. 5) The peak velocity of the initial rise was considered as the highest velocity registered within 200 ms of the rise time. The 6) amplitude and 7) rise time at halfmax were computed at half the CR amplitude. Responses were computed per session and for averages across sessions, only sessions where the CR% exceeded 40% were considered.

### Large-scale electrophysiology using silicon probes

Extracellular recordings were performed using ESSY-37 E1 32 channel silicon probes (Cambridge NeuroTech, UK). An Intan (RHD2132, USA) amplifier was used to digitize and amplify the recorded extracellular voltage signals at 16 bit and were recorded using an Intan RHD2000 Amplifier Evaluation System (sampling rate: 30,000 Hz). We used Open Ephys^76^ for acquisition, online monitoring, and processing of cerebellar electrophysiological signals. A craniotomy 3 mm in diameter was made at AP -6.25 mm, ML -2.25, after which, a cylindrical light-weight recording chamber with a sealable lid was installed at the rim of the craniotomy on the skull surface. During recordings, lidocaine was applied onto the dura surface 15 minutes before recordings, which was subsequently cleaned with saline. For a subset of the mice, a plexiglass grid, designed to fit into the chamber at a horizontal orientation was assembled and the silicon probes were lowered through the grid holes. Our quantification of histological results shows that we recorded from lobules IV/V and Simplex lobules consistently. On a given day, 1-4 sessions were recorded at different depths unidirectionally from ventral to dorsal. The silicon probe was allowed to stabilize for 20 minutes before recording. The mouse could move freely on the treadmill at all times without influencing probe stability.

### Optogenetics

We implanted the tapered optogenetic fiber (Optogenix Sigma fiber) at a 7-degree angle from an adjacent grid hole to cover Purkinje cells recorded from Lobule IV/V and Simplex from a depth of 1125-2000 microns on the ventrolateral aspect. We used mice that expressed ChR2 in Purkinje cells (L7-cre-Ai32), where uniformity of expression was confirmed post hoc through histological analysis of the co-expressing eYFP marker. The power of the LED source was calibrated over pilot experiments to 0.5 mW, which allowed the majority of Purkinje cells recorded to remain below 150 Hz, where the natural firing range of Purkinje cells is 40-200 Hz. Each optogenetics session consisted of 120 trials, on 40% of the trials, a variable pulse train of 20 Hz was delivered to cover the model-predicted time of suppression until after the airpuff. Rasters were sorted based on optogenetic stimulation condition and thereafter on basis of trial condition.

### Unit isolation, Purkinje cell identification and putative molecular layer interneurons

In the absence of optogenetic manipulations, Purkinje cells can also be identified from large-scale in vivo recordings through physiological metrics. We used five criteria for such identification: 1) Recordings were performed from the molecular layer and Purkinje layer based on the polarities of the identified complex spike patterns. 2) The baseline firing rate of neurons lies between 40-200 Hz. 3) Complex spikes and simple spike waveforms were recorded from the same channel or adjacent locations within 20 μm. 4) The complex and simple spike waveforms conformed to standard time scale and shape properties^77^. 5) The complex spike elicited a 20 ms simple spike suppression in a cross-correlogram (with a 10% contamination rate for this criterion). Purkinje cells for which the last condition was not met were labeled putative Purkinje cells. Note that we were unable to record granule cells due to limitations in electrode impedance and data from recording locations in granule layers was excluded when it could be identified. For these reasons, neurons that we could not classify as Purkinje cells are referred to as putative molecular layer interneurons, however, some of these could include cell-types from other layers.

### Functional classification of cerebellar cortical population

Modulatory activity was detected using a recent statistical method known as the ZETA test ^49^. For Purkinje cell simple spiking and putative MLI activity, the test was applied separately to spikes in the paired and test conditions for detecting modulation with respect to the airpuff and the light, respectively. For those neurons showing significant modulations, polarities were determined by comparing baseline and within-trial spiking activity. Baseline spiking was always computed in the -150 to -50 ms window relative to the light. Within-trial spiking for light-modulated activity was computed in test trials in the 100 to 500 ms window following the light. Within-trial spiking for airpuff-modulated activity was computed in airpuff-aligned rasters in the 50 to 200 ms window following the airpuff. For Purkinje cell complex spikes, modulatory activity was detected by applying the ZETA test to windows surrounding events of interest: Light, Prior-onset, and Airpuff. CSpk_Light_ was detected in the 0 to 150 ms window following the light and CSpk_Prior_ was detected in the 190 to 275 ms window following the light, or equivalently, the -10 to 75 ms window surrounding the onset of the prior in the *Wide* condition. Note that the precision of CSpk_Prior_ is much smaller than the full window length. CSpk_Airpuff_ was detected in airpuff-aligned rasters in the 0 to 150 ms window following the airpuff.

### Neural population analysis

We used Principal Component Analysis (PCA) to characterize how the neural population dynamics evolved in each prior condition (*Single*, *Wide*, *Narrow* and *Bimodal* expert conditions and *Single-Wide switch* conditions) as a function of time (Figure 2e,g, Supplementary figure 6a-c). For the *switch* mice, although PCA was applied separately before and after the switch, mice were selected only if there was a comparable number of neurons recorded in both conditions. PCA was then performed on a covariance matrix comprising the collective demeaned firing rates of cerebellar cortical populations (*N* neurons) over time bins *t,* within mouse and within condition *c*, without attrition and with an approximately equal number of trials (see scree plots in Figure 2e, Supplementary Fig. 6a-c).

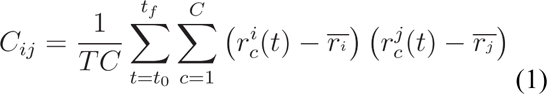

Here *C_ij_* represents the covariance matrix between neurons *i* and *j*, with *T* representing the total number of time bins between *t_f_* and *t*_0_, and *C* the number of conditions within each prior. 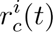 is the average firing rate of neuron *i* under condition *c* at time *t*, and 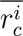 is the mean firing rate of neuron *i* across all time bins, with analogous notation for neuron *j*. The diagonalization of the covariance matrix *C* = *U DU^T^*, yields a new coordinate system given by the columns of the matrix *U*. *D* is a diagonal matrix representing the eigenvalues of the system, indicating the variance captured by the corresponding PCs. The columns of *U* that correspond to the largest eigenvalues, represent the eigenvectors that explain the most variance in the neural data, and are taken in descending order to be the principal components (PC) of the system. The projection of the *N*-dimensional data onto the *k*th PC is given by:

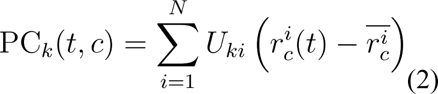

where 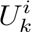 is the *i* element of the *k* PC (*U_k_*). The PCs are linear readouts of the population activity and the contribution of each neuron to a given PC_*k*_(*t*, *c*) is given by the *i*th element of *U_k_*. The first PC accounted for 43% variance on average across mice (further details in Table 2), whereas, the second PC often accounted for less than 10% variance for most subjects (further details in Table 2). We performed a scree analysis to determine the dimensionality for which 80% of the variance could be explained. This number was around 3 or 4 for most mice and conditions. Geometric analysis was therefore performed on test trial projections of the neural population for the first three principal components.

**Table 2:** Percentage of explained variance (%EV) captured by the first two principal components (PC1 & 2) derived from the collective neural activity for each mouse in its corresponding experimental condition.

| %EV Switch |  |  |  | %EV Expert |  |  |  |  |  |
| --- | --- | --- | --- | --- | --- | --- | --- | --- | --- |
| Single |  | Wide |  | Wide |  | Narrow |  | Bimodal |  |
| PC1 | PC2 | PC1 | PC2 | PC1 | PC2 | PC1 | PC2 | PC1 | PC2 |
| 63 | 8 | 63 | 11 | 31 | 13 | 48 | 19 | 57 | 10 |
| 42 | 17 | 42 | 9 | 38 | 14 | 53 | 11 | 31 | 10 |

|  |  |  |  |  |  |  |  |  |  |
| --- | --- | --- | --- | --- | --- | --- | --- | --- | --- |
| 38 | 15 | 38 | 17 | 54 | 10 | 23 | 12 | 42 | 8 |
| 28 | 12 | 28 | 7 | 35 | 11 | 52 | 12 |  |  |
| 46 | 14 | 46 | 14 | 70 | 7 | 45 | 19 |  |  |
| 59 | 13 | 59 | 8 | 44 | 8 | 50 | 8 |  |  |
| 53 | 11 | 53 | 11 | 29 | 7 |  |  |  |  |
| 45 | 13 | 45 | 11 |  |  |  |  |  |  |
| 49 | 18 | 49 | 7 |  |  |  |  |  |  |

### Trial-by-trial analysis of neural activity using LFADS

We used the LFADS framework^51^ to infer latent dynamics from the spike-sorted data recorded in mice during the *Single* expert and *Wide* novice conditions from cerebellar lobule simplex. After successful training runs, LFADS produced a smooth trial-by-trial firing rate estimate for each neuron. We used LFADS in its ‘multi-session stitching’ mode, which enabled the assimilation of multiple datasets originating from the same generative dynamics to be combined in a single neural activity model. We thus combined datasets across sessions but not across prior conditions or mice. We performed hyperparameter tuning of the LFADS models, starting in an apparent underfitting regime at the start of each parameter sweep and incrementally increasing complexity of the factor dimensions and increasing the initial condition bottleneck. The cost function was specified by a Kullback-Leibler (KL), an L2, and reconstruction cost terms. We lowered the starting weight for the KL term before training to prioritize regularization by other means. We also inspected all smooth firing rate inferences and compared those side-by-side with the corresponding spike rasters and peristimulus time histograms (PSTH). We terminated training of networks after their learning rate decreased to below 10^-5^ and we terminated hyperparameter tuning when LFADS networks exhibited sufficient complexity to express the observed condition-averaged firing rates while remaining in the undercomplete regime of the variational autoencoder. We used the LFADS Run Manager and Tensorflow code provided by Pandarinath et al.^51^. The data analysis of the LFADS output was handled through a custom MATLAB-based GUI. Neural data was first filtered to only permit neurons with minimum baseline activity of 5 Hz. Metric extraction on LFADS-inferred traces was performed after removing the mean initial condition variations from each trial. A summary of hyperparameters used for the LFADS models can be found in supplementary table 1. Further, to facilitate the optimization process, we reduced the weight of the KL cost to 0.1 and the batch size for stochastic gradient descent batches to 15 trials.

### Encoding model for time intervals and scalar variability

We assume granule cell spike counts (**r**) obey an inhomogeneous Poisson process whose rate function is Gaussian with mean *ω*(*t*) and standard deviation *σ_i_*. The maximum firing rate of the *i^th^* granule cell parallel fiber, *μ_i_*, is specified with respect to the onset of the light or conditioned stimulus.

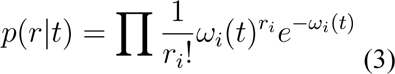

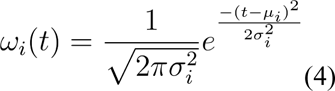

Due to scalar variability, the internal estimate of elapsed time 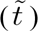 has a probabilistic relationship to the chronometric elapsed time (*t*). We formulated this relationship as a conditional Gaussian probability distribution whose mean is *t* and whose standard deviation remains constant for the *i^th^* kernel but scales across kernels by linear scaling factor *w_b_*, equivalent to the Weber fraction that best describes behavioral observations. Therefore, we assume a heterogeneous population that takes such a form and approximates Weber’s law.

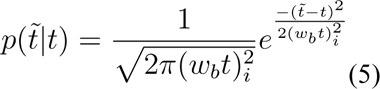

We will now assume a relatively dense and discrete heterogeneous population over stimulus time *t_s_*. Let *p*(*t_s_*) be the prior probability of the stimulus time *t_s_*. While each granule cell may have a preferred firing time, only a subset of granule cells will be active (over elapsed time) when a given *t_s_* is administered. Previous work ^78^ has provided closed-form solutions for infomax allocation of tuning curves as a function of sensory prior knowledge when a heterogeneous basis set if parameterized as follows:

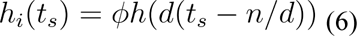

Where, ϕ is a gain term that modulates the maximum average firing rate of each neuron in the population and density *d* controls spacing and width of tuning curves. However, in our case, given the large value of N compared to the timescale of a task, the density term has a lesser impact than in the sensory case described by Ganguli & Simoncelli^78^. On the other hand, if N is too large, the basis set may cease to behave as a low-pass filter. Therefore, there is an optimal value of N for which the model best describes observations in the high noise regime. We assume that for a population of *N* neurons, there is a maximum firing rate **R** that provides an upper bound over the prior and full population:

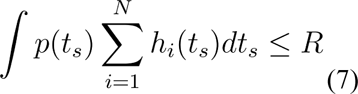

Given these formulations, the Fisher information *I_f_*(*t*) can be written as:

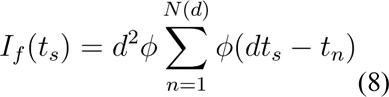

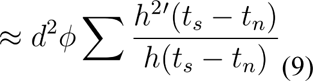

Given the anatomy of the circuit, N is large (we approximate it with 5000 neurons) and we assume densely tiled firing rates based on recent experimental evidence in related tasks^67,68^. This implies that here such a large N is expected to result in a negligible change in density in the post-synaptic effect of the basis set after learning. In other words, after applying realistic anatomical constraints, we find that granule cell tuning may not have to rapidly shift to accommodate the prior. Therefore, the primary factor driving efficient allocation here is modulation to effective firing rates reflecting temporal priors.

Beyond the efficient coding parametrization, the tuning curves of our granule cell population before learning take a specific form to accommodate scalar variability. The firing rate reduction over the population was modeled by a gain function, g(t), with time constant *τ_basis_*, and the increase in width was modeled linearly before learning, *σ_basis_i__*=*σ*_0_*ik*/*N*, where *i* indexes neurons ordered according to their preferred time interval, *N* is the total number of neurons and *k* is the proportion of increase in the width *σ*_0_. The resulting function describes the rate of the *i_th_* granule cell (GC).

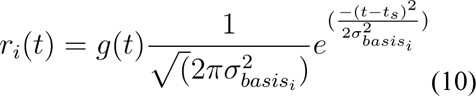

For encoding of *p*(*t_s_*), we define *w_i_*, which represents the postsynaptic weight of the *t_th_* GC with a Purkinje cell. Long-term depression (LTD) in TRACE is modeled for each GC-Purkinje synapse as proportional to the rate of firing of respective GCs shortly before the firing of climbing fibers (CFs) at the time of the airpuff. The time before CF firing at which GC-Purkinje synapses become eligible for LTD is called the eligibility trace (*ϵ*), which we assume occurs 50 ms before the onset of the airpuff in the model. In the absence of CF stimulation and in the presence of GC firing, a weak restoring force (Long-Term Potentiation or LTP) acts to reverse learning. The dynamics of LTD and LTP were governed by their respective time constants, *τ_ltd_* and *τ_ltp_*. In the absence of learning, synapses would gradually drift toward the baseline, *w*_0_.

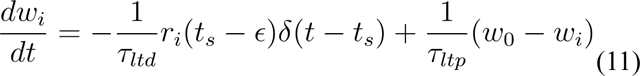

At steady state, the sum of all weights over the basis set population resembles the shape of an inverted prior distribution p(t), which makes sense given that Purkinje cells are inhibitory and one of the primary learning mechanisms in the cerebellar cortex is long-term depression^61^ of Purkinje cell activity. In the model, the change in baseline Purkinje cell activity is computed as a weighted sum of GC activity.

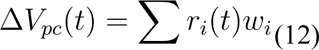

The presence of the eligibility trace implies that any CF firing at the onset of the light cue will have no bearing on the plasticity of the GC-Purkinje cell synapses. Similarly, GC activation at the time of the airpuff will be irrelevant to learning of the prior. Further, our results remain qualitatively unchanged under assumptions of more complex functions of the eligibility trace. Since Purkinje cells constitute the sole output of the cerebellar cortex and disinhibit the cerebellar nuclei (CbN), the primary cerebellar output hypothesized to govern eyeblink behavior, we take the behavioral output to be proportional to

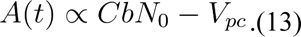

## Author contributions

JK and DN conceived the study. JK, MR and DN performed the experiments. JK, MR, LB, IK, and DN implemented the methods and conducted analyses. All authors wrote the manuscript. DN supervised this project and provided funding for it.

## Acknowledgments

This work was funded by the Netherlands Organization for Scientific Research (Vidi-VI.193.076,

Aspasia-015.016.012 and Gravitation-024.005.022), the AINED organization, and the Nationaal Groeifonds (AINED/NGF NGF.1609.241.021) to DN.

## Conflicts of interest

The authors declare no conflicts of interest exist.

## Data availability statement

All datasets will be made available at an open-access repository like Dryad at the successful conclusion of peer review.

## Code availability statement

Code will be made available at an open-access repository at the successful conclusion of peer review.

## Supplementary Figures

**Supplementary figure 1:**
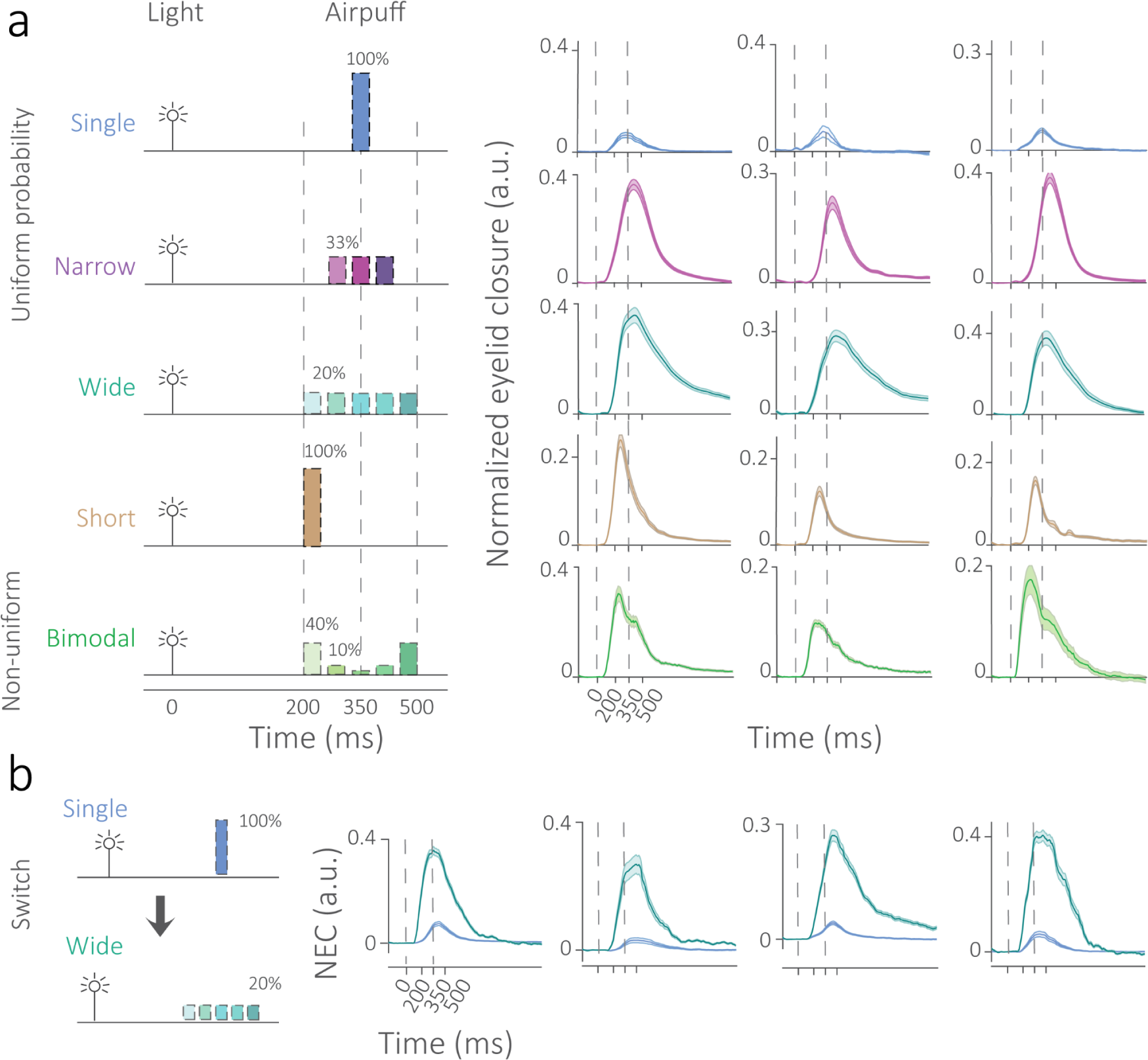
Behavioral traces on test trials. a) Examples of average behavioral traces from individual subjects on test trials (airpuff is omitted) for the *Single* (blue), *Narrow* (purple), *Wide* (teal), *Short* (brown), *Bimodal* (green) prior conditions. Solid lines represent averages, shaded areas represent standard error of the mean. b) Same subject switched from *Single* (blue) to *Wide* (teal) prior condition. Average eyelid closure traces for four subjects. Solid lines represent averages, shaded areas represent standard error of the mean.

**Supplementary figure 2:**
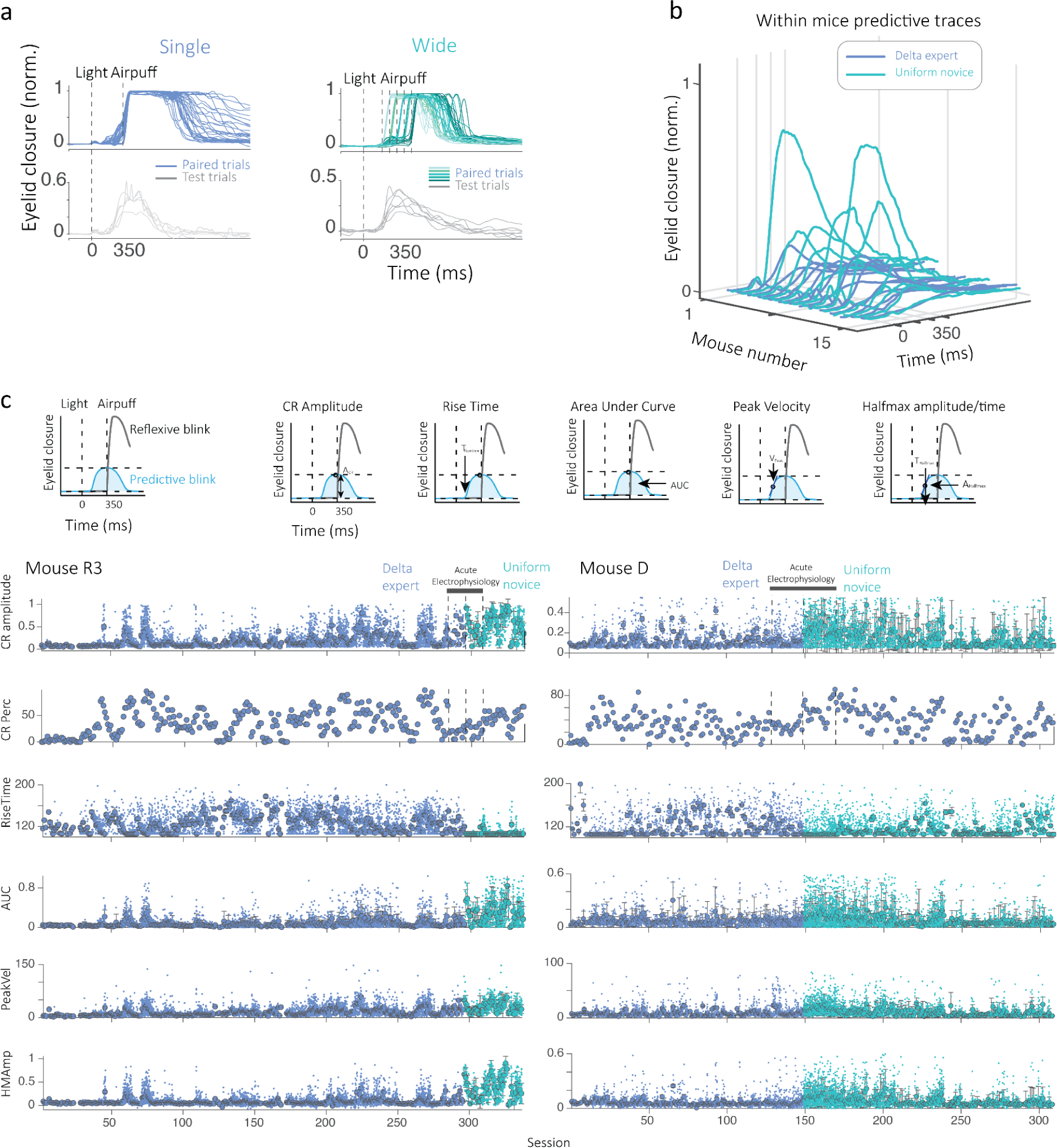
Eyelid behavior of *switch* mice across sessions. Normalized trial-wise eyelid closure traces for *Single* expert paired (blue) and test trials (gray) and *Wide* expert paired (teal) and test trials (gray). b) Test traces for eyelid closure of all mice (N = 15) that transitioned from the *Single* (blue) to *Wide* (teal) condition. c) Above: Description of various metrics used to quantify the predictive eyeblink. Below: Session-wise progression of learning metrics for two mice that transitioned from *Single* expert to *Wide* novice. The period where electrophysiology was performed is also indicated.

**Supplementary figure 3:**
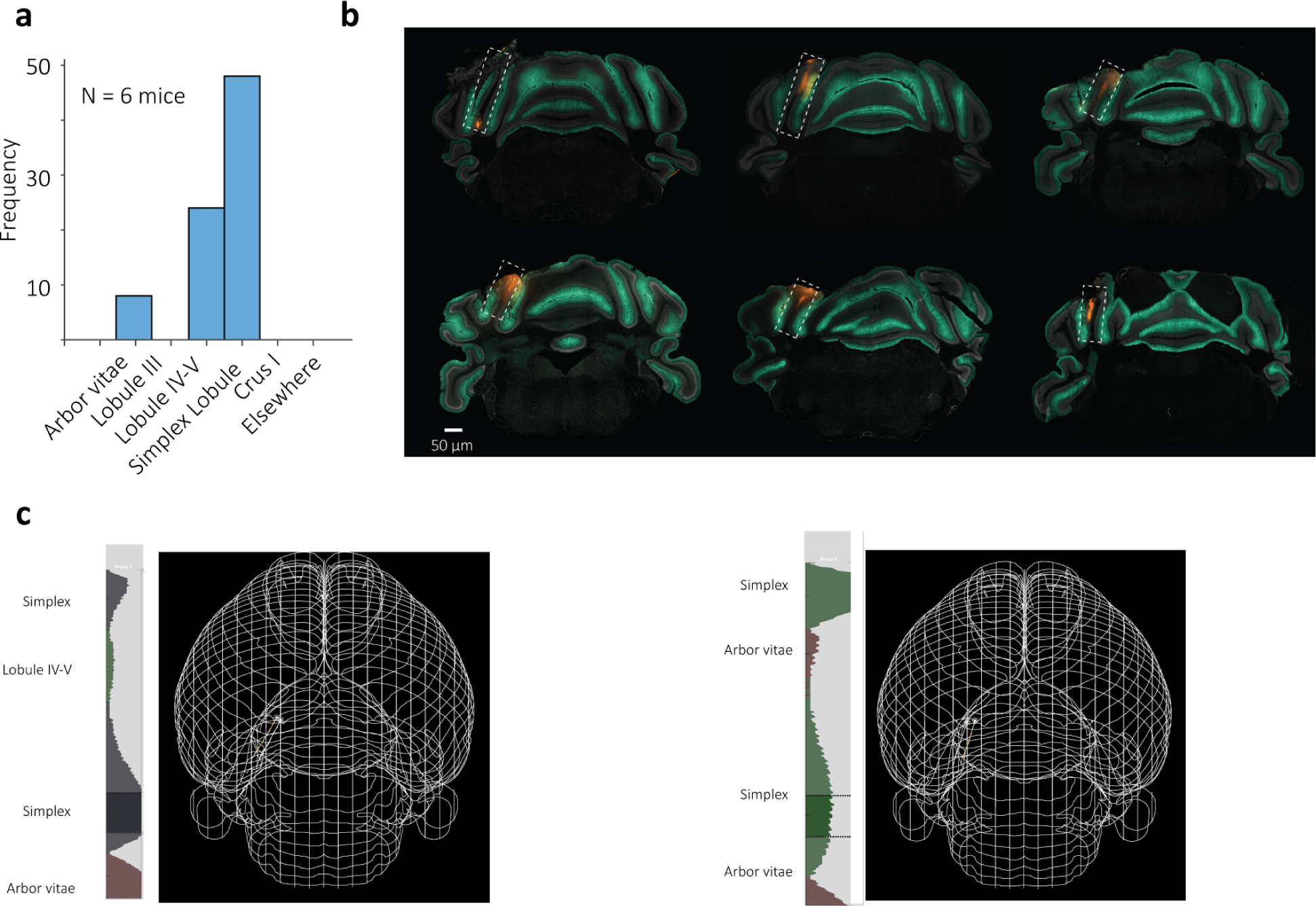
Quantification of recording sites. a) Histogram of locations of detected probe track points aligned to Allen Common coordinate framework (CCF). b) Examples of histological images used to identify probe entry locations as marked with DiI at grid point epicenter using a single shank on the last day of recording. The entry point consistently lies between lobules Simplex and IV/V. Recordings were made from ventral (max depth - 2200 um) to dorsal locations (max depth 300 um) with nonlinear double-shank electrodes. Note that perfusion was not performed on the same day after track-marking since animals continued to train on the *Wide* condition for several days for behavioral analysis. c) Allen CCF probe track analysis examples and probe tracks marked in a 3D mesh after registration with the Allen CCF framework for two mice.

**Supplementary figure 4:**
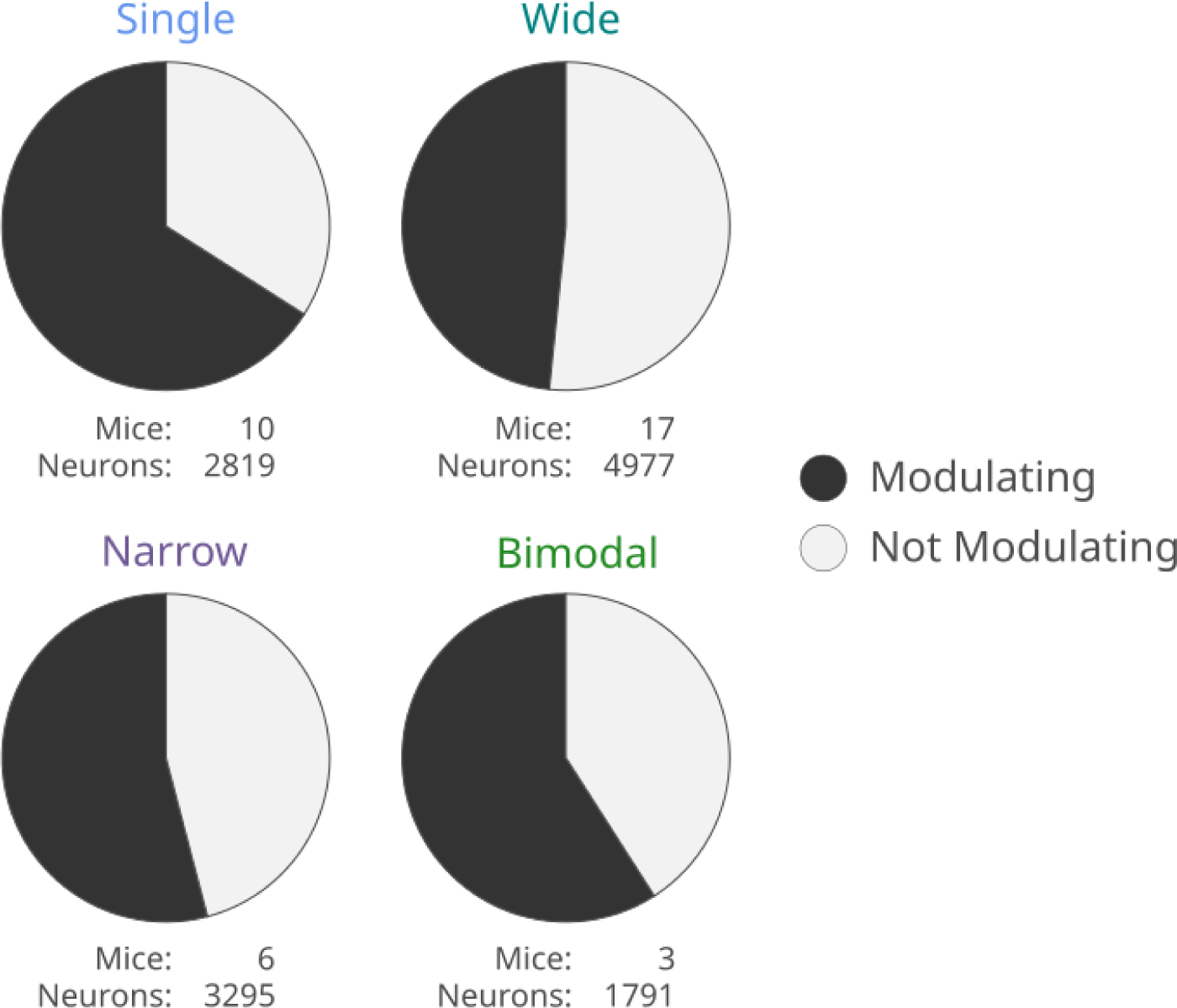
Counts from statistical test that determined task-related modulation. Top left to bottom right: Pie charts showing percentage and numbers of modulating (black) and non-modulating (white) neurons for the *Single*, *Wide*, *Narrow*, and *Bimodal* conditions.

**Supplementary figure 5:**
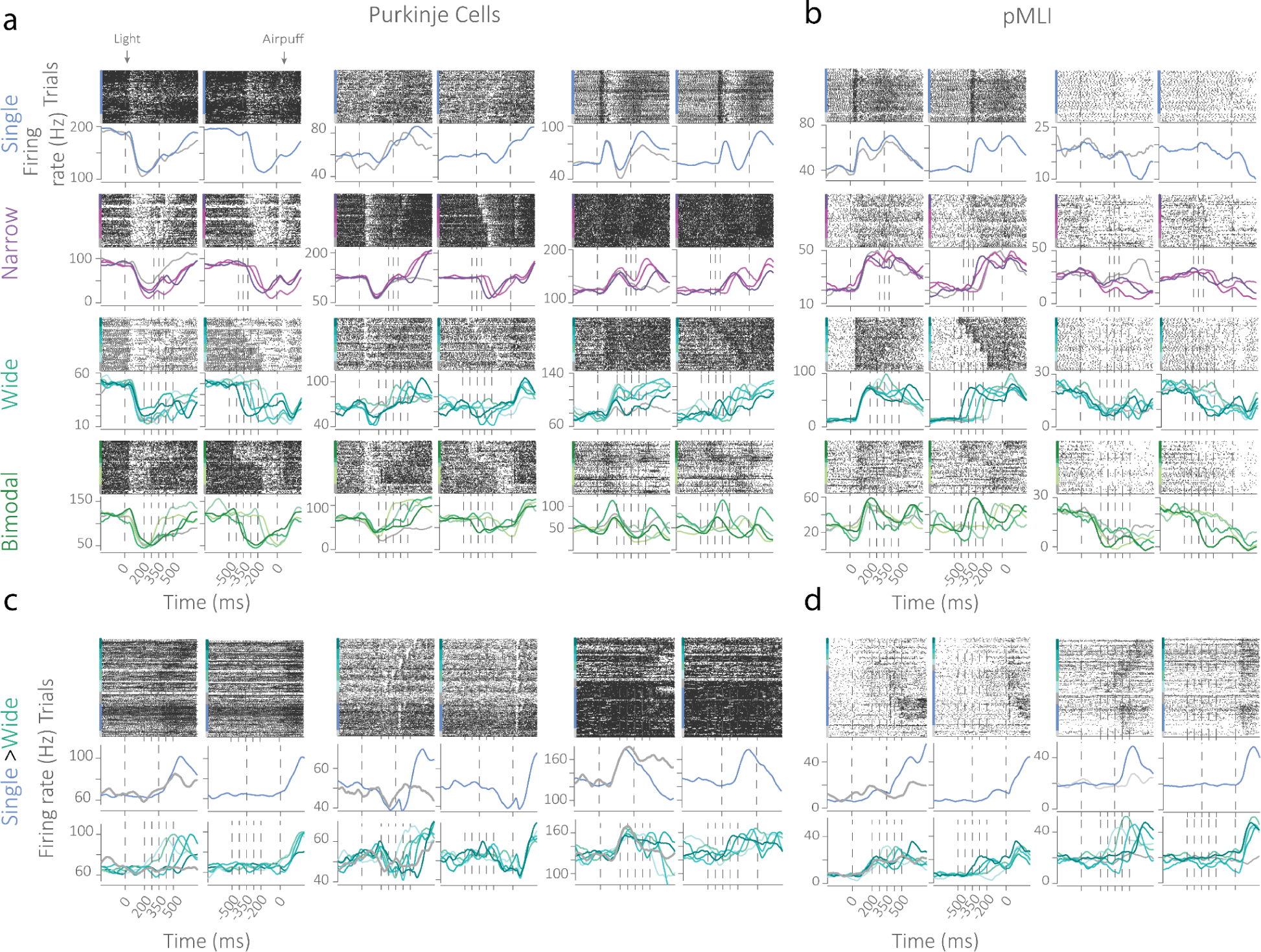
Examples of single units in various conditions. a) Purkinje cell rasters and firing rates over time, aligned to onset of the light (left panel) and airpuff (right panel) for the *Single* (blue), *Narrow* (purple), *Wide* (teal), and *Bimodal* (green) conditions. b) Same as a) for putative molecular layer interneurons (pMLI) for the different prior conditions. c) Purkinje cell rasters and firing rates over time for *switch* mice from the last *Single* session (blue) to the first *Wide* session (teal). d) Same as c) for pMLIs.

**Supplementary figure 6:**
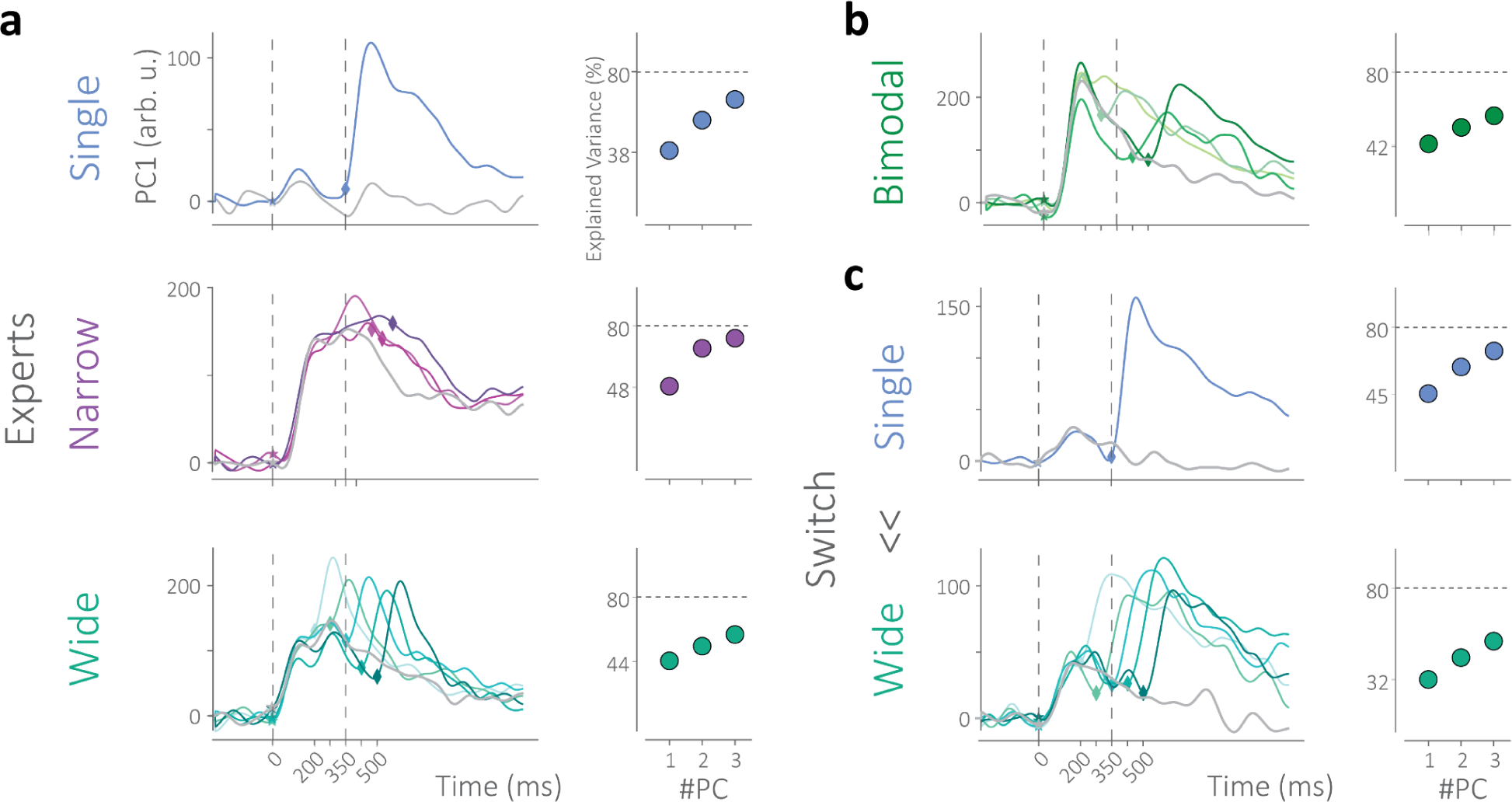
Principal components recapitulate behavioral results. a) Left: Examples of within-individual evolution of the first principal component of neural population activity for paired and test conditions for the *Single, Narrow* and *Wide* conditions. Right: Variance explained (%) of the first three principal components for each individual and condition. b) Same as a) for the *Bimodal* prior condition. c) Same as a for the *switch* condition as the mouse experienced the last *Single* session and switched to the first *Wide* session.

**Supplementary figure 7:**
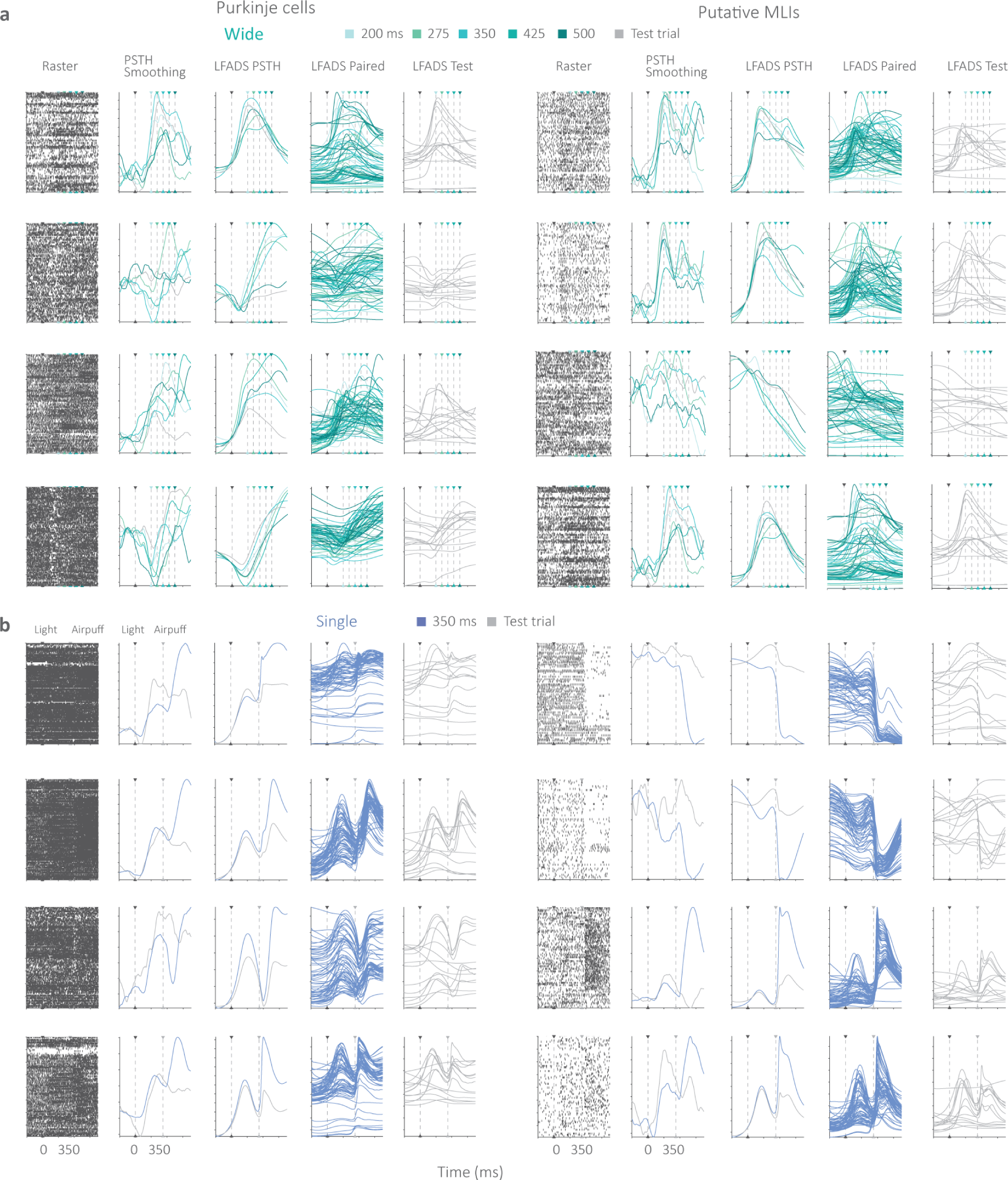
Examples of LFADS estimates. a) Left to right: Spike times organized trial-wise (raster), conditioned average firing rates, historically known as peri-stimulus time histograms or PSTH, conditioned averaged LFADS-inferred activity, LFADS inference of activity on individual trials in the *Wide* condition. Left: examples of Purkinje cells, Right: examples of putative molecular layer interneurons. b) Same as a) for the *Single* condition.

**Supplementary figure 8:**
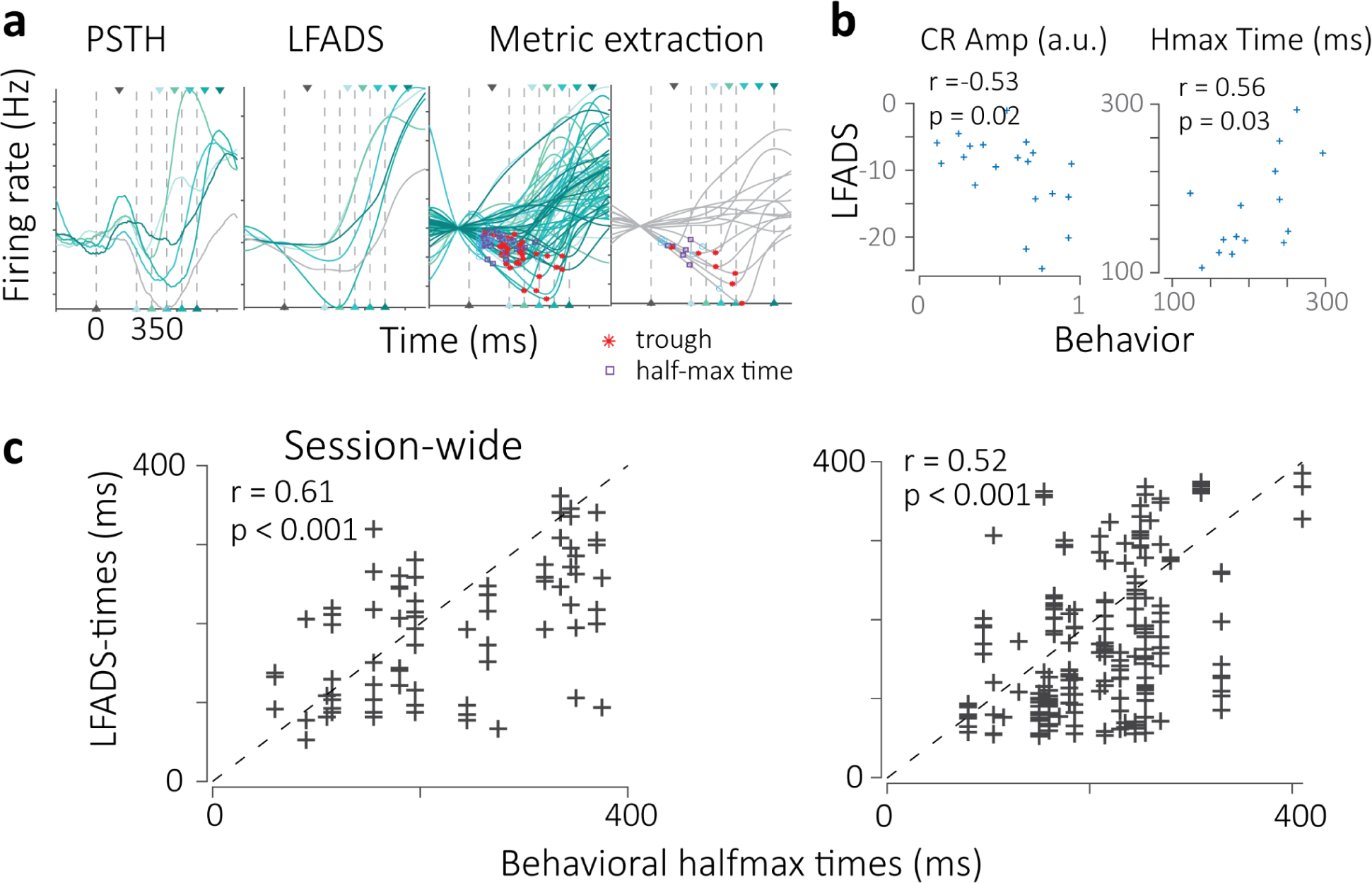
LFADS trial-by-trial metric decoding. a) Comparison of condition-averaged firing rates of neural data vs. condition-averaged LFADS-estimates of firing rates and individual trialwise estimates for the paired and test conditions. The trough (minima, asterisk) and half-max time (square) estimates are indicated. b) Correlation of behaviorally extracted metrics of CR-amplitude and half-max time to LFADS estimates of firing rate trough and half-max time. c) Session-wide comparison between behavioral metrics and LFADS-derived metrics for half-max time.

**Supplementary figure 9:**
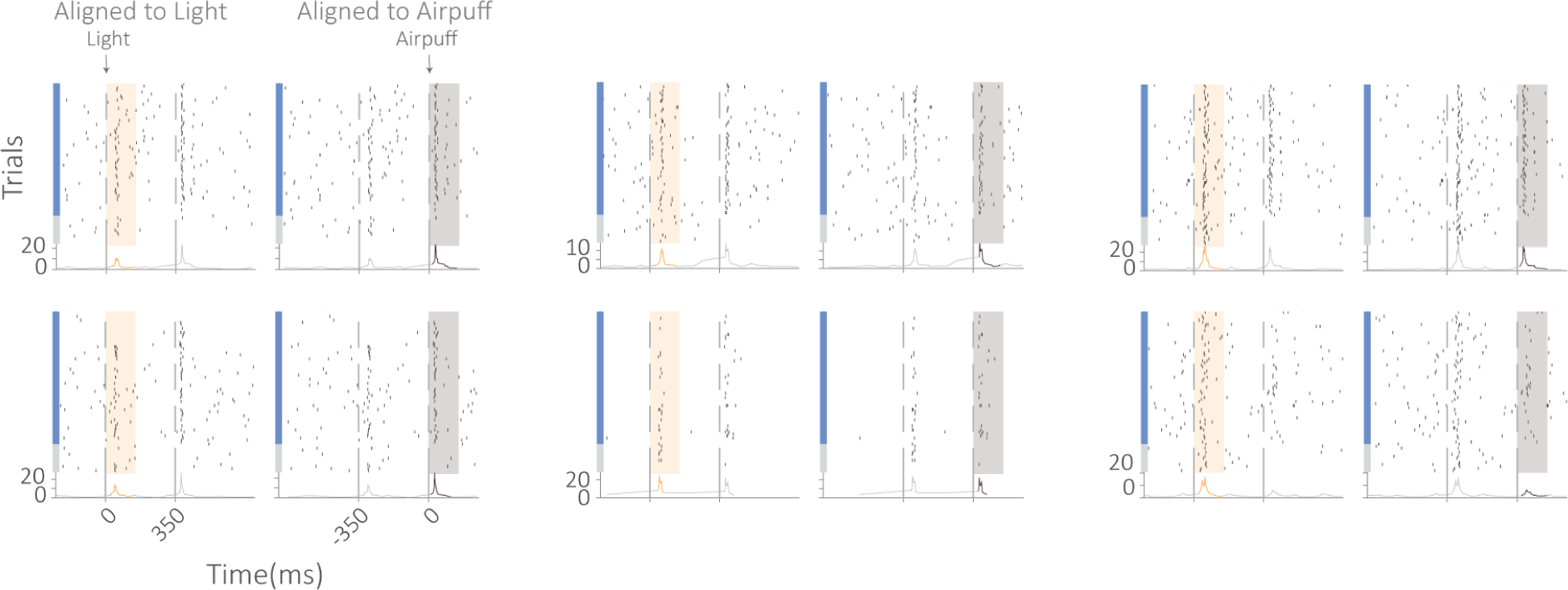
Purkinje cell complex spikes in the *Single* condition. Top: Complex spike rasters aligned to the light (left panel) and airpuff (right panel). Bottom: Instantaneous firing rate estimates over time inferred from multiple timescales of spiking activity ^49^. The CSpk_light_ is shown in orange and the CSpk_airpuff_ in gray.

**Supplementary figure 10:**
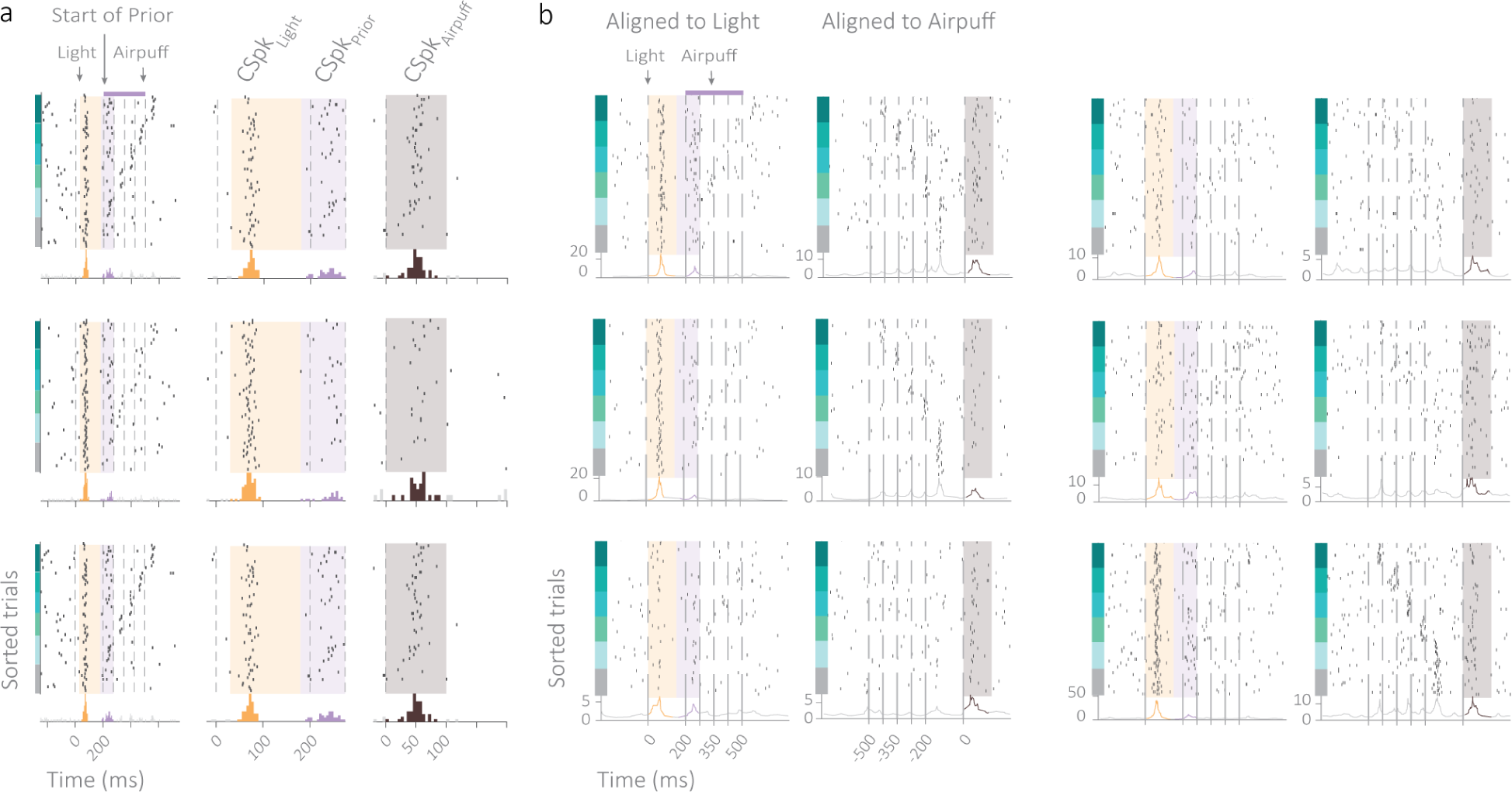
Purkinje cell complex spikes in the *Wide* condition. a) In addition to CSpk_Light_ and CSpk_Airpuff_ signals, we additionally encounter a complex spike signal following the onset of the prior, CSpk_Prior_. b) Top: Complex spike rasters aligned to the light (left panel) and airpuff (right panel). Bottom: Instantaneous firing rate estimates over time inferred from multiple timescales of spiking activity ^49^. The light, prior, and airpuff signals are shown in orange, purple, and gray, respectively.

**Supplementary figure 11:**
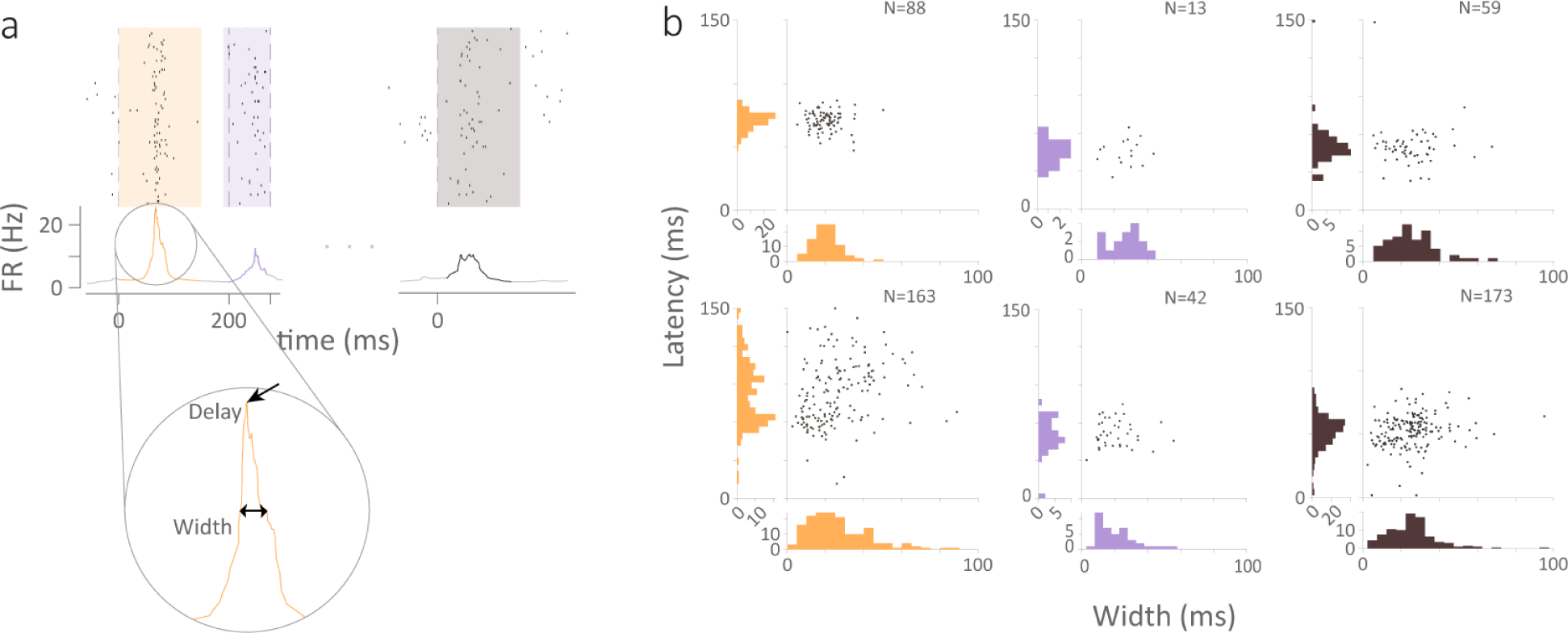
Characterization of the width and latency of complex spike signals. a) Each delay is computed as the amount of time following the event after which the firing rate peaks. Width is computed as the amount of time that passes between the two points where the firing rate attains half of the peak amplitude. b) Examples of widths vs. delays computed for all CSpks in a given mouse. CSpk_Light_, CSpk_Prior_, and CSpk_Airpuff_ are shown in orange, purple, and black, respectively.

**Supplementary figure 12:**
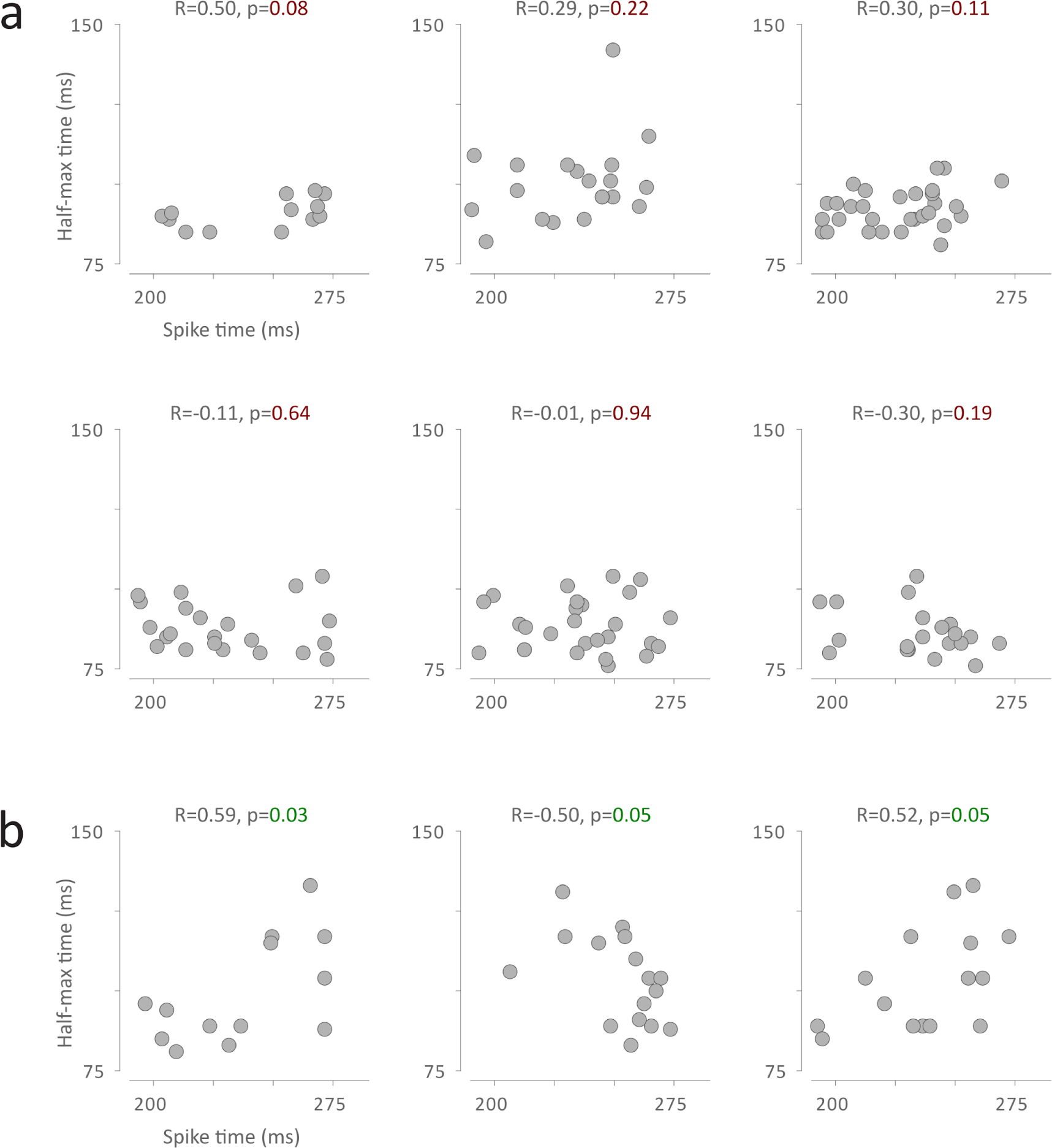
a) Examples of non-significant trial-by-trial correlation of behavioral times and the time of CSpk_prior_. b) same as a) for significant examples of correlation. Vast majority of cells did not show any statistically significant correlation with behavioral metrics.

**Supplementary figure 13:**
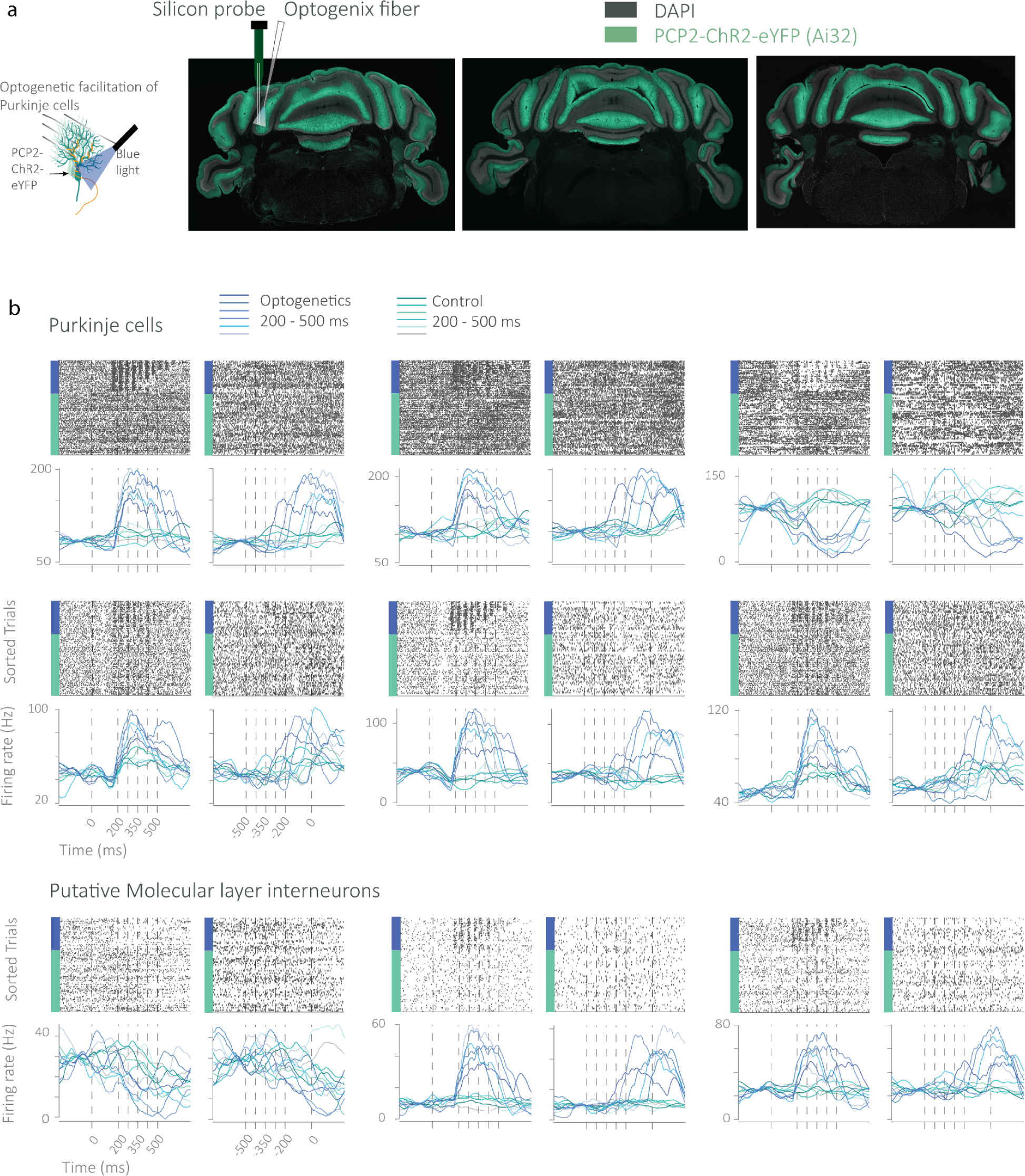
Expression of ChR2 and electrophysiology during optogenetics. a) Left: Schematic of optogenetic strategy to acutely stimulate Purkinje cells expressing Channelrhodopsin2 (ChR2) with blue light. Three coronal cross-sections of Ai32 mutants show uniform expression of ChR2 in Purkinje cells (L7xAi32). Electrophysiological recordings were made from lobule simplex where an Optogenix tapered fiber was also implanted. b) Purkinje cell and putative MLI rasters and firing rates over time in the *Wide* condition during optogenetic trials and control trials within the same session. Optogenetic perturbation was applied at random on 40% of the trials but the rasters have been sorted for visualization.

## Supplementary Tables

**Supplementary Table 1:** Hyperparameters used to train LFADS models in *Single* and *Wide* conditions.

| LFADS hyperparameters | Single | Wide |
| --- | --- | --- |
| spikeBinMs | 1 | 1 |
| c_co_dim | 0 | 0 |
| c_batch_size | 15 | 15 |
| c_factors_dim | 40 | 40 |
| c_gen_dim | 64 | 64 |
| c_ic_enc_dim | 64 | 64 |
| c_learning_rate_stop | 1.00E-05 | 1.00E-05 |
| c_kl_ic_weight | 0.1 | 0.1 |
| c_ic_dim | 50 | 50 |
| c_learning_rate_n_to_compare | 3 | 3 |
| useAlignmentMatrix | TRUE | TRUE |

**Supplementary Table 2:** Neuron counts per prior condition.

|  | <u>Single</u> | <u>Narrow</u> | <u>Wide Novice</u> | <u>Wide Expert</u> | <u>Bimodal</u> |
| --- | --- | --- | --- | --- | --- |
| *No. Mice* | 10 | 6 | 9 | 8 | 3 |
| *No. Neurons* | 2819 | 3295 | 2638 | 2339 | 1791 |
| *No. Modulating* | 1860 | 1782 | 1342 | 1066 | 1058 |
| *% Modulating* | 65.98 | 54.08 | 50.87 | 45.58 | 59.07 |

## Notes

### Competing Interest Statement

The authors have declared no competing interest.

## References

1. Körding, K. P. & Wolpert, D. M. Bayesian integration in sensorimotor learning. Nature 427, 244–247 (2004).

2. Jazayeri, M. & Shadlen, M. N. Temporal context calibrates interval timing. Nat. Neurosci. 13, 1020–1026 (2010).

3. Ma, W. J. & Jazayeri, M. Neural coding of uncertainty and probability. Annu. Rev. Neurosci. 37, 205–220 (2014).

4. Knill, D. C. & Richards, W. Perception as Bayesian Inference. (Cambridge University Press, 1996).

5. Knill, D. C. & Pouget, A. The Bayesian brain: the role of uncertainty in neural coding and computation. Trends Neurosci. 27, 712–719 (2004).

6. Doya, K., Ishii, S. & Pouget, A. Bayesian Brain: Probabilistic Approaches to Neural Coding. (MIT Press, 2007).

7. Rao, R. P. N., Olshausen, B. A. & Lewicki, M. S. Probabilistic Models of the Brain: Perception and Neural Function. (MIT Press, 2002).

8. Körding, K. P. & Wolpert, D. M. Bayesian decision theory in sensorimotor control. Trends Cogn. Sci. 10, 319–326 (2006).

9. Mamassian, P. & Landy, M. S. Interaction of visual prior constraints. Vision Res. 41, 2653–2668 (2001).

10. Adams, W. J., Graf, E. W. & Ernst, M. O. Experience can change the ‘light-from-above’ prior. Nat. Neurosci. 7, 1057–1058 (2004).

11. Stocker, A. A. & Simoncelli, E. P. Noise characteristics and prior expectations in human visual speed perception. Nat. Neurosci. 9, 578–585 (2006).

12. Jacoby, N. & McDermott, J. H. Integer Ratio Priors on Musical Rhythm Revealed Cross-culturally by Iterated Reproduction. Curr. Biol. 27, 359–370 (2017).

13. Girshick, A. R., Landy, M. S. & Simoncelli, E. P. Cardinal rules: visual orientation perception reflects knowledge of environmental statistics. Nat. Neurosci. 14, 926–932 (2011).

14. Acerbi, L., Wolpert, D. M. & Vijayakumar, S. Internal representations of temporal statistics and feedback calibrate motor-sensory interval timing. PLoS Comput. Biol. 8, e1002771 (2012).

15. Norton, E. H., Acerbi, L., Ma, W. J. & Landy, M. S. Human online adaptation to changes in prior probability. PLoS Comput. Biol. 15, e1006681 (2019).

16. Berniker, M., Voss, M. & Kording, K. Learning priors for Bayesian computations in the nervous system. PLoS One 5, (2010).

17. Dehaene, G. P., Coen-Cagli, R. & Pouget, A. Investigating the representation of uncertainty in neuronal circuits. PLoS Comput. Biol. 17, e1008138 (2021).

18. Ma, W. J., Beck, J. M., Latham, P. E. & Pouget, A. Bayesian inference with probabilistic population codes. Nat. Neurosci. 9, 1432–1438 (2006).

19. Fiser, J. & Lengyel, G. A common probabilistic framework for perceptual and statistical learning. Curr. Opin. Neurobiol. 58, 218–228 (2019).

20. Darlington, T. R., Beck, J. M. & Lisberger, S. G. Neural implementation of Bayesian inference in a sensorimotor behavior. Nat. Neurosci. 21, 1442–1451 (2018).

21. Sohn, H., Narain, D., Meirhaeghe, N. & Jazayeri, M. Bayesian Computation through Cortical Latent Dynamics. Neuron 103, 934–947.e5 (2019).

22. Miyazaki, M., Nozaki, D. & Nakajima, Y. Testing Bayesian models of human coincidence timing. J. Neurophysiol. 94, 395–399 (2005).

23. Li, Y. & Dudman, J. T. Mice infer probabilistic models for timing. Proc. Natl. Acad. Sci. U. S. A. 110, 17154–17159 (2013).

24. Ohmae, S., Uematsu, A. & Tanaka, M. Temporally specific sensory signals for the detection of stimulus omission in the primate deep cerebellar nuclei. J. Neurosci. 33, 15432–15441 (2013).

25. Uematsu, A., Ohmae, S. & Tanaka, M. Facilitation of temporal prediction by electrical stimulation to the primate cerebellar nuclei. Neuroscience 346, 190–196 (2017).

26. ten Brinke, M. M. et al. Evolving Models of Pavlovian Conditioning: Cerebellar Cortical Dynamics in Awake Behaving Mice. Cell Rep. 13, 1977–1988 (2015).

27. Heiney, S. A., Wohl, M. P., Chettih, S. N., Ruffolo, L. I. & Medina, J. F. Cerebellar-dependent expression of motor learning during eyeblink conditioning in head-fixed mice. J. Neurosci. 34, 14845–14853 (2014).

28. Ten Brinke, M. M. et al. Dynamic modulation of activity in cerebellar nuclei neurons during pavlovian eyeblink conditioning in mice. Elife 6, (2017).

29. Albergaria, C., Silva, N. T., Pritchett, D. L. & Carey, M. R. Locomotor activity modulates associative learning in mouse cerebellum. Nat. Neurosci. 21, 725–735 (2018).

30. Becker, M. I. & Person, A. L. Cerebellar Control of Reach Kinematics for Endpoint Precision. Neuron 103, 335–348.e5 (2019).

31. Guo, J.-Z. et al. Disrupting cortico-cerebellar communication impairs dexterity. Elife 10, (2021).

32. Tsutsumi, S. et al. Purkinje Cell Activity Determines the Timing of Sensory-Evoked Motor Initiation. Cell Rep. 33, 108537 (2020).

33. Halverson, H. E., Khilkevich, A. & Mauk, M. D. Cerebellar Processing Common to Delay and Trace Eyelid Conditioning. J. Neurosci. 38, 7221–7236 (2018).

34. Khilkevich, A., Zambrano, J., Richards, M.-M. & Mauk, M. D. Cerebellar implementation of movement sequences through feedback. Elife 7, (2018).

35. Kalmbach, B. E. et al. Cerebellar cortex contributions to the expression and timing of conditioned eyelid responses. J. Neurophysiol. 103, 2039–2049 (2010).

36. Schlerf, J. E., Spencer, R. M. C., Zelaznik, H. N. & Ivry, R. B. Timing of rhythmic movements in patients with cerebellar degeneration. Cerebellum 6, 221–231 (2007).

37. Ivry, R. B. & Keele, S. W. Timing functions of the cerebellum. J. Cogn. Neurosci. 1, 136–152 (1989).

38. Chettih, S. N., McDougle, S. D., Ruffolo, L. I. & Medina, J. F. Adaptive timing of motor output in the mouse: the role of movement oscillations in eyelid conditioning. Front. Integr. Neurosci. 5, 72 (2011).

39. Ito, M. Control of mental activities by internal models in the cerebellum. Nat. Rev. Neurosci. 9, 304–313 (2008).

40. Wolpert, D. M., Chris Miall, R. & Kawato, M. Internal models in the cerebellum. Trends in Cognitive Sciences 2, 338–347 (1998).

41. Imamizu, H. et al. Human cerebellar activity reflecting an acquired internal model of a new tool. Nature 403, 192–195 (2000).

42. Lisberger, S. G. Internal models of eye movement in the floccular complex of the monkey cerebellum. Neuroscience 162, 763–776 (2009).

43. Brooks, J. X., Carriot, J. & Cullen, K. E. Learning to expect the unexpected: rapid updating in primate cerebellum during voluntary self-motion. Nat. Neurosci. 18, 1310–1317 (2015).

44. Laurens, J. & Angelaki, D. E. How the Vestibulocerebellum Builds an Internal Model of Self-motion. The Neuronal Codes of the Cerebellum 97–115 (2016).

45. Laurens, J., Meng, H. & Angelaki, D. E. Neural representation of orientation relative to gravity in the macaque cerebellum. Neuron 80, 1508–1518 (2013).

46. Mackrous, I., Carriot, J., Jamali, M. & Cullen, K. E. Cerebellar Prediction of the Dynamic Sensory Consequences of Gravity. Curr. Biol. 29, 2698–2710.e4 (2019).

47. Raymond, J. L. & Medina, J. F. Computational Principles of Supervised Learning in the Cerebellum. Annu. Rev. Neurosci. 41, 233–253 (2018).

48. Wang, Q. et al. The Allen Mouse Brain Common Coordinate Framework: A 3D Reference Atlas. Cell 181, 936–953.e20 (2020).

49. Montijn, J. S. et al. A parameter-free statistical test for neuronal responsiveness. Elife 10, (2021).

50. De Zeeuw, C. I., Koppen, J., Bregman, G. G., Runge, M. & Narain, D. Heterogeneous encoding of temporal stimuli in the cerebellar cortex. Nat. Commun. 14, 7581 (2023).

51. Pandarinath, C. et al. Inferring single-trial neural population dynamics using sequential auto-encoders. Nat. Methods 15, 805–815 (2018).

52. Ohmae, S. & Medina, J. F. Climbing fibers encode a temporal-difference prediction error during cerebellar learning in mice. Nat. Neurosci. 18, 1798–1803 (2015).

53. Mauk, M. D., Steinmetz, J. E. & Thompson, R. F. Classical conditioning using stimulation of the inferior olive as the unconditioned stimulus. Proc. Natl. Acad. Sci. U. S. A. 83, 5349–5353 (1986).

54. McCormick, D. A., Steinmetz, J. E. & Thompson, R. F. Lesions of the inferior olivary complex cause extinction of the classically conditioned eyeblink response. Brain Res. 359, 120–130 (1985).

55. Heiney, S. A., Kim, J., Augustine, G. J. & Medina, J. F. Precise control of movement kinematics by optogenetic inhibition of Purkinje cell activity. J. Neurosci. 34, 2321–2330 (2014).

56. Madisen, L. et al. A toolbox of Cre-dependent optogenetic transgenic mice for light-induced activation and silencing. Nat. Neurosci. 15, 793–802 (2012).

57. Wang, X., Yu, S.-Y., Ren, Z., De Zeeuw, C. I. & Gao, Z. A FN-MdV pathway and its role in cerebellar multimodular control of sensorimotor behavior. Nat. Commun. 11, 6050 (2020).

58. Yeo, C. H. & Hesslow, G. Cerebellum and conditioned reflexes. Trends Cogn. Sci. 2, 322–330 (1998).

59. Marr, D. A theory of cerebellar cortex. J. Physiol. 202, 437–470 (1969).

60. Albus, J. S. A theory of cerebellar function. Mathematical Biosciences 10 25–61 (1971).

61. Ito, M. & Kano, M. Long-lasting depression of parallel fiber-Purkinje cell transmission induced by conjunctive stimulation of parallel fibers and climbing fibers in the cerebellar cortex. Neurosci. Lett. 33, 253–258 (1982).

62. Medina, J. F. & Mauk, M. D. Computer simulation of cerebellar information processing. Nat. Neurosci. 3 Suppl, 1205–1211 (2000).

63. Yamazaki, T. & Tanaka, S. Computational models of timing mechanisms in the cerebellar granular layer. Cerebellum 8, 423–432 (2009).

64. Narain, D., Remington, E. D., Zeeuw, C. I. D. & Jazayeri, M. A cerebellar mechanism for learning prior distributions of time intervals. Nat. Commun. 9, 469 (2018).

65. Garcia-Garcia, M. G. et al. A cerebellar granule cell-climbing fiber computation to learn to track long time intervals. Neuron (2024) doi:10.1016/j.neuron.2024.05.019.

66. Kennedy, A. et al. A temporal basis for predicting the sensory consequences of motor commands in an electric fish. Nat. Neurosci. 17, 416–422 (2014).

67. Giovannucci, A. et al. Cerebellar granule cells acquire a widespread predictive feedback signal during motor learning. Nat. Neurosci. 20, 727–734 (2017).

68. Wagner, M. J., Kim, T. H., Savall, J., Schnitzer, M. J. & Luo, L. Cerebellar granule cells encode the expectation of reward. Nature 544 96–100 (2017).

69. Fujita, M. Adaptive filter model of the cerebellum. Biol. Cybern. 45, 195–206 (1982).

70. Barri, A., Wiechert, M. T., Jazayeri, M. & DiGregorio, D. A. Synaptic basis of a sub-second representation of time in a neural circuit model. Nat. Commun. 13, 7902 (2022).

71. Mauk, M. D. & Ruiz, B. P. Learning-dependent timing of Pavlovian eyelid responses: differential conditioning using multiple interstimulus intervals. Behav. Neurosci. 106, 666–681 (1992).

72. Bowers, J. S. & Davis, C. J. Bayesian just-so stories in psychology and neuroscience. Psychol. Bull. 138, 389–414 (2012).

73. Griffiths, T. L., Chater, N., Norris, D. & Pouget, A. How the Bayesians got their beliefs (and what those beliefs actually are): comment on Bowers and Davis (2012). Psychological bulletin 138 415–422 (2012).

74. Zhang, X.-M. et al. Highly restricted expression of Cre recombinase in cerebellar Purkinje cells. Genesis 40, 45–51 (2004).

75. Heiney, S. A., Ohmae, S., Kim, O. A. & Medina, J. F. Single-Unit Extracellular Recording from the Cerebellum During Eyeblink Conditioning in Head-Fixed Mice. Neuromethods 134, 39–71 (2018).

76. Siegle, J. H. et al. Open Ephys: an open-source, plugin-based platform for multichannel electrophysiology. J. Neural Eng. 14, 045003 (2017).

77. Kostadinov, D., Beau, M., Blanco-Pozo, M. & Häusser, M. Predictive and reactive reward signals conveyed by climbing fiber inputs to cerebellar Purkinje cells. Nat. Neurosci. 22, 950–962 (2019).

78. Ganguli, D. & Simoncelli, E. P. Efficient sensory encoding and Bayesian inference with heterogeneous neural populations. Neural Comput. 26, 2103–2134 (2014).

